# Meta-ecosystem dynamics drive the spatial distribution of functional groups in river networks

**DOI:** 10.1101/2021.06.04.447105

**Authors:** Claire Jacquet, Luca Carraro, Florian Altermatt

## Abstract

The meta-ecosystem concept provides a theoretical framework to study the effect of local and regional flows of resources on ecosystem dynamics. Meta-ecosystem theory has hitherto been applied to highly abstract landscapes, and meta-ecosystem dynamics in real-world landscapes remain largely unexplored. River networks constitute a prime example of meta-ecosystems, being characterized by directional resource flows from upstream to downstream communities and from the terrestrial to the aquatic realm. These flows have been thoroughly described by the River Continuum Concept (RCC), a seminal concept in freshwater ecology, stating that observed spatial variations in the relative abundances of invertebrate functional groups reflect systematic shifts in types and locations of food resources, which are in turn determined by the physical attributes of river reaches. Hence, the RCC represents a solid conceptual basis for determining how changes in landscape structure and resource flows will translate into local and regional changes in community composition. Here, we develop and analyse a riverine meta-ecosystem model inspired by the RCC, which builds upon a physically-based landscape model of dendritic river networks. We show that the spatial distributions and regional biomass of invertebrate functional groups observed in stream communities are determined by the spatial structure and scaling attributes of dendritic river networks, as well as by specific rates of resource flows. Neglecting any of these aspects in modelling river meta-ecosystems would result in unrealistic community patterns. Moreover, we observed that high rates of resource flow, for example due to river anthropization, have a negative effect on the regional biomass of all functional groups studied, and can lead to cascading extinctions at the meta-ecosystem scale. Our work paves the way for the development of physically-based meta-ecosystem models to understand the structure and functioning of real-world ecosystems.

## Introduction

Ecosystems are open to flows of materials, organisms, and energy, and understanding the effects of these flows on ecosystem structure and functioning is a central goal of ecological research (Polis et al., 2004). By integrating the local production and movement of abiotic resources into metacommunity theory, the meta-ecosystem framework allows investigating feedback processes between community and resource dynamics across spatial scales (Gounand et al., 2018a; Loreau et al., 2003; Massol et al., 2011; Guichard and Marleau, 2021). Seminal works on meta-ecosystems have been developed on simplified landscapes (Loreau et al., 2003; Loreau and Holt, 2004), often represented by two-patch ecosystems (Gravel et al., 2010a,b; Marleau et al., 2010). This simple representation allowed an analytical investigation of the interplay between local and regional flows of matter and their implications for community dynamics (Gravel et al., 2010a; Loreau et al.,2003; Leroux and Loreau, 2012; Massol et al., 2011). More recent studies focusing on larger spatial networks outlined the importance of spatial structure and movement rates of organisms and materials to promote meta-ecosystem stability (Marleau et al., 2014; Gravel et al., 2016). However, one enduring limitation of current meta-ecosystem models is the abstract representation of the landscape (Gounand et al., 2018a), which is often described by random or Cartesian spatial networks (Marleau et al., 2014; Gravel et al., 2016). In contrast, the physical structure of real-world landscapes constrains organisms’ movement and resource flows and is likely to influence the spatial distributions of resources and the organisms that feed upon them (Harvey et al., 2017a, 2020; Leroux and Loreau, 2008; Montagano et al., 2018; Schmitz et al., 2018).

River networks constitute a prime example of meta-ecosystems, as documented by a large body of literature in freshwater ecology assessing the critical influence of directional resource flows, both from the surrounding terrestrial environment and from upstream river reaches, on the composition of local stream communities (e.g., Soininen et al. (2015); Abelho and Descals (2019); Bartels et al. (2012)). Evidence that composition of local riverine communities is linked with the river system in its entirety was prominently pointed out in the River Continuum Concept (RCC) (Vannote et al., 1980), a cornerstone concept in freshwater ecology. The RCC proposes that commonly observed spatial variations in the relative abundances of major functional groups of organisms along a longitudinal river gradient reflect systematic shifts in types and locations of food resources, which are in turn determined by the physical attributes of river reaches (e.g., stream width). The organismal groups considered are freshwater invertebrates that are generally clustered in five functional groups (grazers, shredders, collectors, filterers and predators), which are of high relevance with respect to the biodiversity and functioning of riverine ecosystems (Anderson and Sedell, 1979; Wallace et al., 2015; Harvey and Altermatt, 2019). The RCC specifically predicts that shredders should be the most abundant functional group in small river reaches, which are strongly shaded by the surrounding vegetation and receive a large input of dead organic matter from falling leaves. The abundance of grazers is expected to peak at mid-sized streams, where light penetration into the stream is highest, stimulating the development of primary producers. Light penetration, and therefore primary production, is limited by water depth and turbidity in large rivers, reducing the abundance of grazers. Finally, filterers and collectors should be the most abundant groups in large rivers where the most abundant resources is fine particulate organic matter, which is a by-product of leaf consumption by shredders and is delivered into the water column from upstream communities.

The RCC is likely the most influencing conceptual framework in freshwater and stream ecology and is among the most commonly cited works in this field (totalling >6000 citations in the Scopus database), prompting empirical and theoretical works assessing its strengths and limitations (see Doretto et al. (2020) for a recent review). Although the parallel between meta-ecosystem theory and the mechanisms formulated in the RCC has been acknowledged in several recent studies (Massol et al., 2011; Doretto et al., 2020; Gounand et al., 2018a; Harvey et al., 2020), a formal integration of the RCC within a meta-ecosystem model is still lacking. Next to the explicit predictions mentioned above, the RCC further implicitly predicts that changes in the physical attributes of a river network should translate into local and regional changes in the composition of invertebrate communities. However, this prediction has never been investigated formally and it remains unclear how changes in resource input and transport will affect the spatial distribution of invertebrate functional groups in a river network. Furthermore, the RCC essentially describes river systems as a linear array of sites, hence disregarding the contribution of river dendritic structure on the spatial patterns observed in river meta-ecosystems (Doretto et al., 2020).

In this study, we develop a spatially explicit meta-ecosystem model for river systems inspired by the RCC to investigate the effect of meta-ecosystem dynamics on the functional composition of stream communities. We specifically address the following research questions: (i) What is the contribution of the dendritic structure of river networks on the spatial distribution of functional groups described in the RCC? (ii) How do changes in resource flow rate influence the spatial distribution and regional biomass of functional groups in river networks? To do so, we make use of a physically-based model of dendritic river networks, which expresses how stream characteristics (e.g., water discharge, stream width) vary across a river system as a function of drainage area (i.e., the portion of land over which precipitation is drained towards a given river cross-section).

We compare the spatial distributions of functional groups between a dendritic river network and a linear river channel and demonstrate that the spatial patterns described in the RCC can only emerge from a meta-ecosystem model that accounts for the dendritic structure of river networks. We then analyse different scenarios of resource spatial dynamics and show that increased rates of resource flows have a negative impact on the regional biomass of all the functional groups studied and can lead to extinctions at the meta-ecosystem scale.

## Methods

### Meta-ecosystem structure and dynamics

We considered a riverine meta-ecosystem composed of a set of local ecosystems, each of them defined by a river reach and its surrounding terrestrial area, which are spatially connected via resource flow across the river network. A river reach corresponds to an uninterrupted stretch of river in which abiotic conditions can be assumed as constant, and hence constitutes the fundamental unit of a river network. In our model, each local ecosystem is composed of four abiotic resources, a group of primary producers and five groups of consumers, corresponding to the trophic groups and resource types commonly found in freshwater ecosystems (Doretto et al., 2020; Larsen et al., 2019; Vannote et al.,1980; Wallace and Webster, 1996).

First, grazers *G* feed on primary producers *P* (algae or aquatic rooted vascular plants), which are supported by nutrients *N* (e.g., nitrogen) and light (Fig. 1). Second, three independent functional groups feed on different forms of particulate organic matter derived from decomposed terrestrial leaf litter: shredders *S*, collectors *C*, and filterers *F* feed on coarse (*CPOM*), fine (*FPOM*), and dissolved (*DOM*) particulate organic matter, respectively. Finally, predators *R* feed on other groups of consumers, that is *G*, *S*, *C* and *F* (Fig. 1). The abiotic resources present in a local ecosystem originate both from local terrestrial inputs of *N*, *CPOM* and DOM and from upstream reaches via hydrological transport of *N*, *CPOM*, *FPOM* and *DOM*. Consequently, local ecosystems are connected through hydrological transport of resources, and the composition of functional groups in upstream ecosystems determines not only the local resource consumption, but also the amount of resources available for the ecosystems situated downstream. Note that Vannote et al. (1980) originally included both filterers and collectors in the same functional group (i.e., collectors), while a distinction between collectors (feeding on *FPOM*) and filterers (feeding on *DOM*) has been operated in more recent literature (see e.g. Larsen et al., 2019; Doretto et al., 2020). Hence, the densities of collectors described in Vannote et al. (1980) should be compared with the sum of collector and filterer densities in our model.

**Figure 1:**
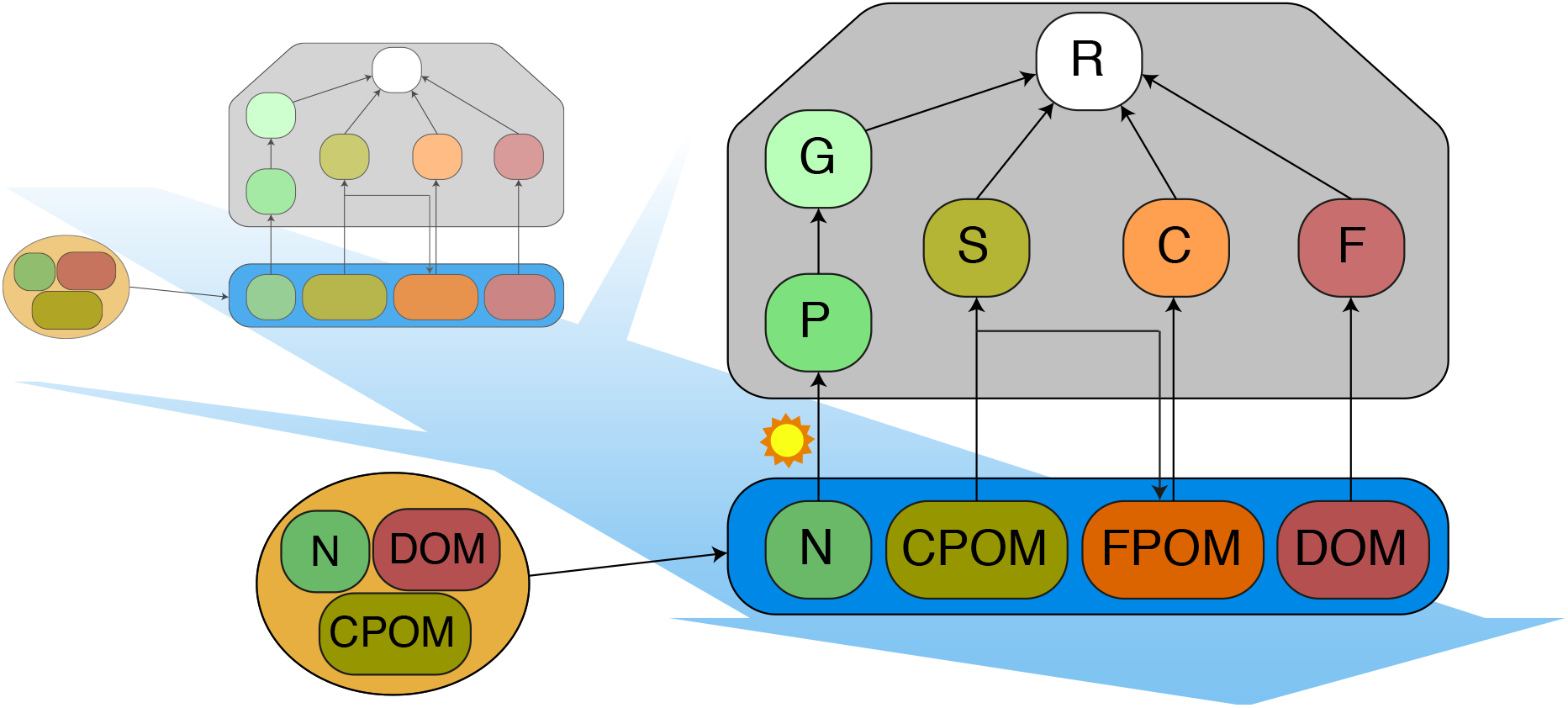
Conceptual illustration of the meta-ecosystem model for river networks. A local stream community (grey box) is composed of a primary producer *P* and five functional groups of consumers (*G*, *S*, *C*, *F* and *R*). The black arrows illustrate the feeding links between the groups and the abiotic resources (blue box). The resources available for a given local community originate both from local terrestrial inputs (yellow ellipse on the left) and from the ecosystems situated upstream via hydrological transport (blue arrow and transparent boxes). Dissolved nutrients (*N*) and light are the basal resources for primary producers *P*, while coarse (*CPOM*), fine (*FPOM*) and dissolved (*DOM*) particulate organic matter are the basal resources for shredders (*S*), collectors (*C*) and filterers (*F*), respectively. Predators (*R*) can feed on all primary consumers, that is *G*, *S*, *C* and *F*. By feeding on *CPOM*, shredders (*S*) produce *FPOM* that constitutes the main resource for collectors (*C*).

We focused our analysis on an equilibrium state for the meta-ecosystem, and did not consider the temporal fluctuation of resource inputs, such as seasonality in resource availability or stream flow. We further hypothesized that living organisms only move within river reaches and do not disperse across local ecosystems. Our meta-ecosystem model is expressed by the following set of ordinary differential equations:

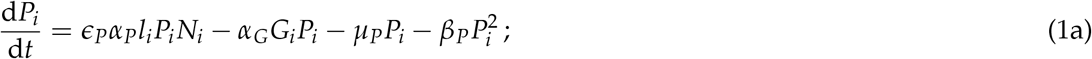

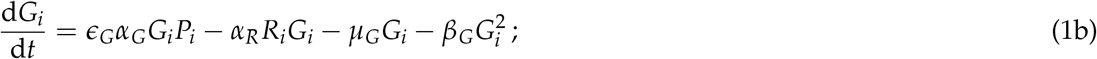

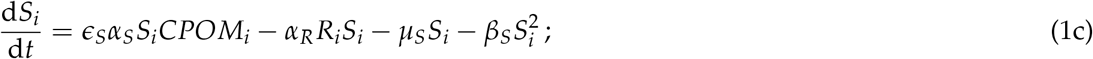

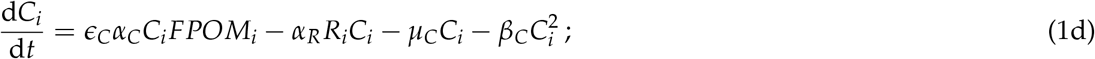

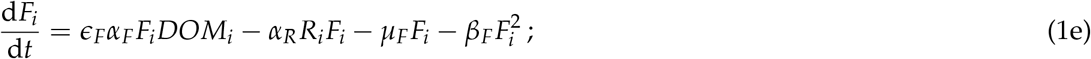

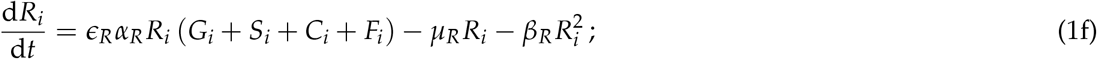

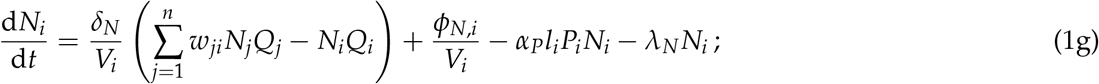

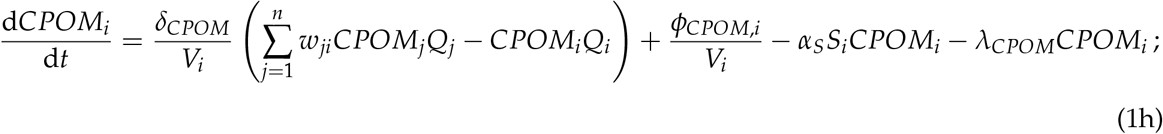

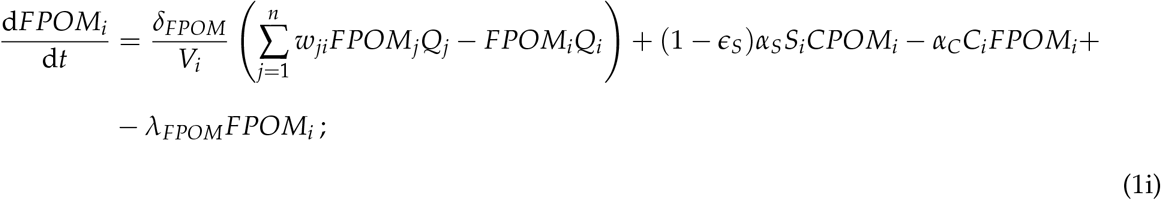

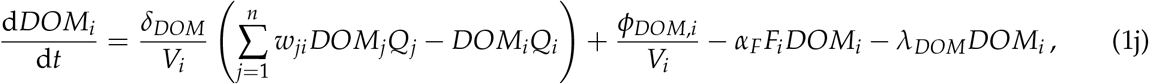

where subscript *i* identifies a river reach. System Eq. (1) is based on a Lotka-Volterra formulation of trophic interactions (including self regulation terms, following Barbier and Loreau (2019)), and on mass balance equations (Eqs. (1g)–(1j)) for the dynamics of resources in river networks. For the living compartments *X* = {*P*, *G*, *S*, *C*, *F*, *R*}(Eqs. (1a–f)), the rate of change in biomass density *X* depends on biomass gain owing to feeding (assuming direct dependence of feeding rate on biomass production) and biomass loss due to metabolism, predation and intra-group competition for resources (density dependence). The parameters related to the living compartments *X* are feeding rate *α_X_*, efficiency of resource assimilation *ε_X_*, strength of intra-group competition *β_X_*, and mortality rate *μ_X_*.

To include the effect of light availability on nutrient uptake of producers, we expressed the realized assimilation rate of producers as *_P_l_i_*, where *l_i_* is a site-specific light limitation factor derived from physical principles and by following the downwelling irradiance concept (Fasham et al., 1990; Davies-Colley and Nagels, 2008). To derive *l_i_*, we assumed that the irradiance of photosynthetically active radiation above the canopy is constant across the river system, while spatial variations in *l_i_* are solely determined by variations in river geometry (i.e., width and depth–see Supporting Information for details). Where available, parameters were chosen in agreement with established evidence on feeding behaviour for the various functional groups. Detailed information on parameter values is available in Table S1.

For the resource compartments *ϒ* ={*N*, *CPOM*, *FPOM*, *DOM*} (Eqs. (1g–j)), the rate of change of resource density *ϒ* depends on mass gain from local terrestrial inputs and inputs from upstream reaches via hydrological transport; and mass loss due to consumption by living organisms, downstream hydrological transport, and resource degradation and deposition. The parameters related to the abiotic resources *ϒ* are the flux of local terrestrial inputs *φ_ϒ,i_*, the relative downstream velocity of resource *ϒ* with respect to water *δ_ϒ_* (i.e., if *δ_ϒ_* = 0.5, resource *ϒ* travels downstream half as fast as water), and the rate of resource loss due to processes other than consumption *λ_ϒ_* (i.e., degradation or deposition). *Q_i_* and *V_i_* represent the water discharge and the water volume of reach *i*, respectively; *w_ji_* is the generic entry of the adjacency matrix (*w_ji_* = 1 if river reach *j* drains into *i* and 0 otherwise); *n* is the total number of reaches constituting the river network.

Following Vannote et al. (1980); Marks (2019), we assumed that *FPOM* is a by-product of *CPOM* consumption by shredders (Fig. 1), therefore inputs of *FPOM* originate from the aquatic environment only. Importantly, Eqs. (1g–j) outline how the physical attributes of river networks, that is water discharge *Q_i_* and volume *V_i_*, influence the density of resources available locally for the functional groups. In particular, both water discharge and volume determine the velocity at which resources are transported across a given reach, while water volume also influences the concentration of local terrestrial inputs.

### Resource spatial dynamics in river networks

We generated a large, virtual river network (so-called optimal channel network, OCN) with the R-package *OCNet* (Carraro et al., 2020, 2021) in order to study changes in the abundance density of functional groups along a gradient of physical conditions specific to river networks. OCNs are structures that reproduce the topological connectivity and scaling features of real river networks (Rinaldo et al., 2014) and are well suited to study riverine ecological processes (Carraro et al., 2020). We built the OCN (Fig. 2a) on a square lattice spanning an area of 5625 km^2^ partitioned into 3688 reaches in order to obtain a gradient of physical conditions sufficiently large to reproduce the predictions of the RCC, which apply to river systems spanning a wide range of stream sizes (see details in Supporting Information).

**Figure 2:**
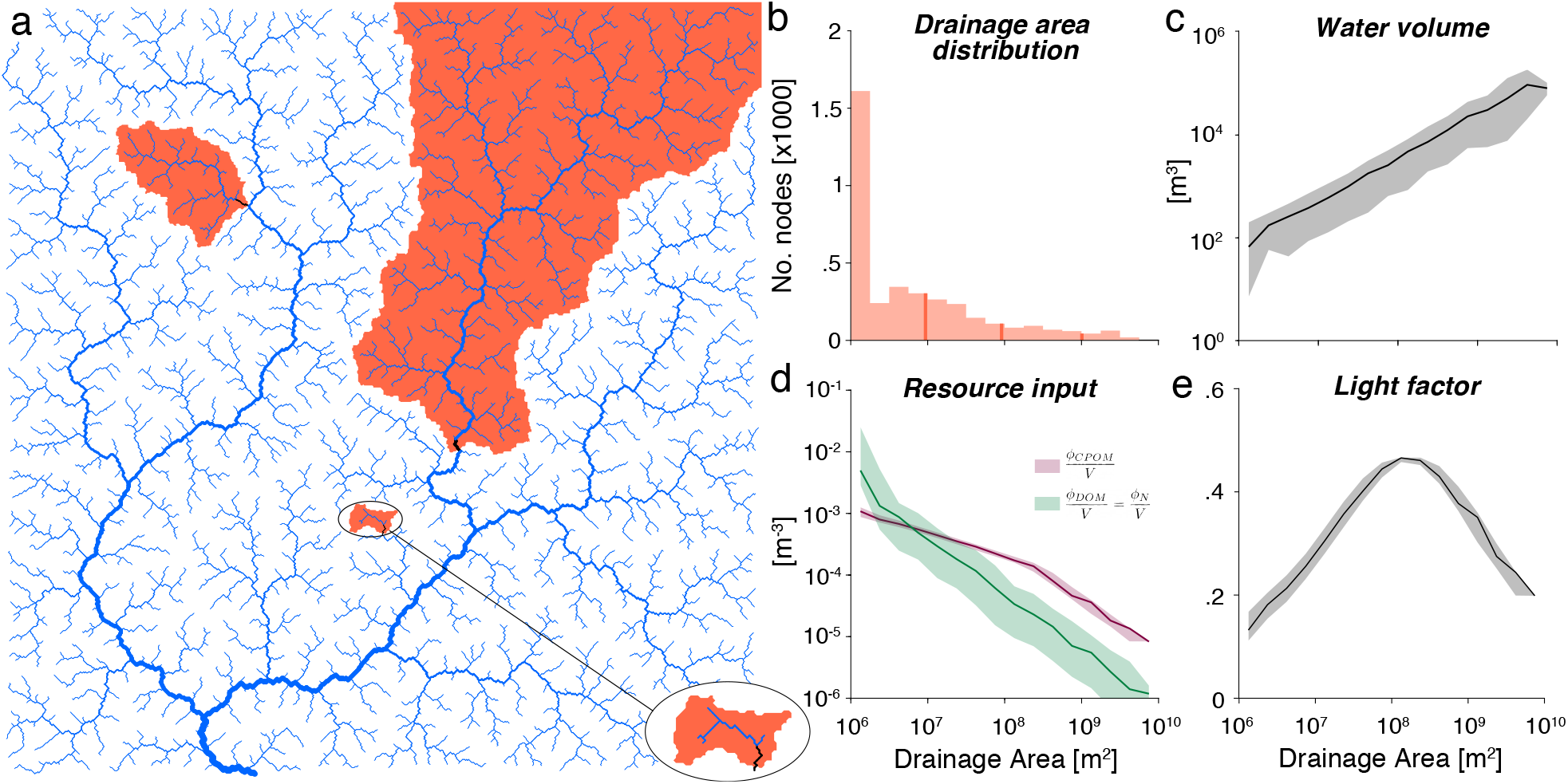
a) The dendritic river network used in the model simulation, spanning a square of area 5625 km^2^. The river network is partitioned into 3688 reaches. Three example reaches are highlighted in black, and their corresponding drainage areas (i.e., the portion of land over which water drains into a given reach) are shown in orange; a zoom-in for the smallest of these reaches is provided at the bottom-right corner. b) Distribution of drainage area values across the 3688 reaches constituting the river network, ranging from 1 to 5625 km^2^ (grouped into 16 bins). Orange solid lines display the drainage area values corresponding to the three orange areas illustrated in panel a. c–e) Variation of physical attributes of the river network variables along a gradient of drainage area: c) water volume increases with drainage area, d) concentration of terrestrial inputs decreases with drainage area, and e) light availability is maximum for intermediate values of drainage area (around 100 km^2^). In panels c–e, solid lines represent median values calculated over reaches corresponding to each bin of panel b, while shaded areas correspond to the 2.5^th^–97.5^th^ percentile intervals evaluated across the bins of panel b.

We made several assumptions regarding the rate of hydrological transport of resources in order to incorporate the verbal arguments of the RCC into our meta-ecosystem model. In particular, we assumed *CPOM* to be transported downstream at a low rate (*δ_CPOM_* = 0.01) compared to water because of the large size of its constituents, which likely induces clogging (Vannote et al., 1980; Wallace and Webster, 1996). Conversely, we assumed *FPOM* to travel downstream at an intermediate rate (*δ_FPOM_* = 0.5), while *DOM* and *N* are transported at a high rate (i.e., same velocity as water: *δ_DOM_* = 1 and *δ_N_* = 1) (Cushing et al., 1993; Wallace and Webster, 1996).

We derived water discharge *Q_i_* and water volume *V_i_* (Fig. 2c) across all river reaches of the network based on drainage area values (Fig. 2b) via the scaling relationships of Leopold and Maddock (1953). Drainage area corresponds to the portion of land over which precipitation is drained towards a given location (Fig. 2a). As a universal geomorphological feature, drainage area is the master variable controlling the physical and hydrological characteristics of a given reach (Leopold et al., 1964; Rodriguez-Iturbe and Rinaldo, 2001). We therefore used drainage area to describe the positioning of a reach within the river network and illustrated how water volume (Fig. 2c), concentration of terrestrial inputs (Fig. 2e), and light availability (Fig. 2e) change along a gradient of drainage area (details on how these variables are calculated are reported in the Supporting Information). Specifically, the concentration of all resources decreases as drainage area increases (Fig. 2d) because of the corresponding increase in water volume along the downstream direction (Fig. 2c). Note that local inputs of *CPOM* depend on river width and not on drainage area, therefore *CPOM* follows a slightly different pattern than other resources in Fig. 2d (see Supporting Information for more details). Light availability *l_i_* peaks at intermediate values of drainage area (Fig. 2e) and is lower both in upstream (due to increased shading effect of canopy in narrower reaches) and downstream reaches (due to increased water depth and subsequent limited light penetration).

We determined the equilibrium densities of each compartment of the meta-ecosystem in all reaches of the river network by finding a feasible equilibrium state (i.e., non-negative densities for all state variables (including resources) at all reaches) for system Eq. (1) by using a linearization method (see Supporting Information for details). We investigated how the densities of functional groups change along a gradient of drainage area and compared the model predictions with the empirical patterns described in the River Continuum Concept (Vannote et al., 1980).

### Alternative scenarios of landscape structure and resource dynamics

We assessed the influence of assuming a complex, dendritic landscape on the spatial distribution of functional groups by running the meta-ecosystem model (Eq. (1)) on an alternative landscape based on minimalist assumptions, namely a linear river channel (without branches). We maintained the following quantities equal in the two landscapes: number of reaches, total drainage area, water discharge at the outlet, total river length, total water volume, and total resource inputs from the terrestrial realm. We further imposed river width, depth and water volume to be constant along the linear river channel and independent from drainage area (see details in Supporting Information). Consequently, light availability is constant as well and does not peak at intermediate values of drainage area in the linear river channel. We compared the spatial distributions of functional groups between the original dendritic network and the linear river channel. Additionally, in the Supporting Information we provide details on the effect of different assumptions in the formulation of linear and dendritic landscapes with respect to the scaling of physical variables.

For the dendritic landscape, we further compared the spatial distributions and regional biomass of each functional group for two alternative scenarios of resource spatial dynamics: a “no flow” scenario, where hydrological transport of resources is neglected, and a “fast” scenario where all resources are transported at a high rate (i.e., same velocity as water). All the scenarios were based on the same set of parameters specified in Table S1.

## Results

The meta-ecosystem model for dendritic river networks accurately reproduces the distributions of functional groups described in the River Continuum Concept (Fig. 3). Specifically, the density of grazers peaks at intermediate values of drainage area (Fig. 3a,f), which is mirrored by the analogous pattern for primary producers (Fig. 3g). Both of these patterns essentially follow the spatial distribution of light availability (Fig. 2e). The density of shredders is maximum in headwaters and decreases as drainage area increases (Fig. 3b,f), while filterers tend to be more homogeneously distributed across the river network (Fig. 3d,f). Conversely, the density of collectors monotonically increases with drainage area (Fig. 3c,f). Consequently, collectors and filterers are the most abundant groups in largest river reaches. In 468 reaches (12.7% of all reaches), all located at the headwaters, collectors are predicted to go extinct. Finally, the density of predators (Fig. 3e), which is proportional to the sum of their preys’ densities (Fig. 3f), is relatively constant along a gradient of drainage area. The corresponding spatial patterns of resource densities predicted by the model are shown in Fig. S1.

**Figure 3:**
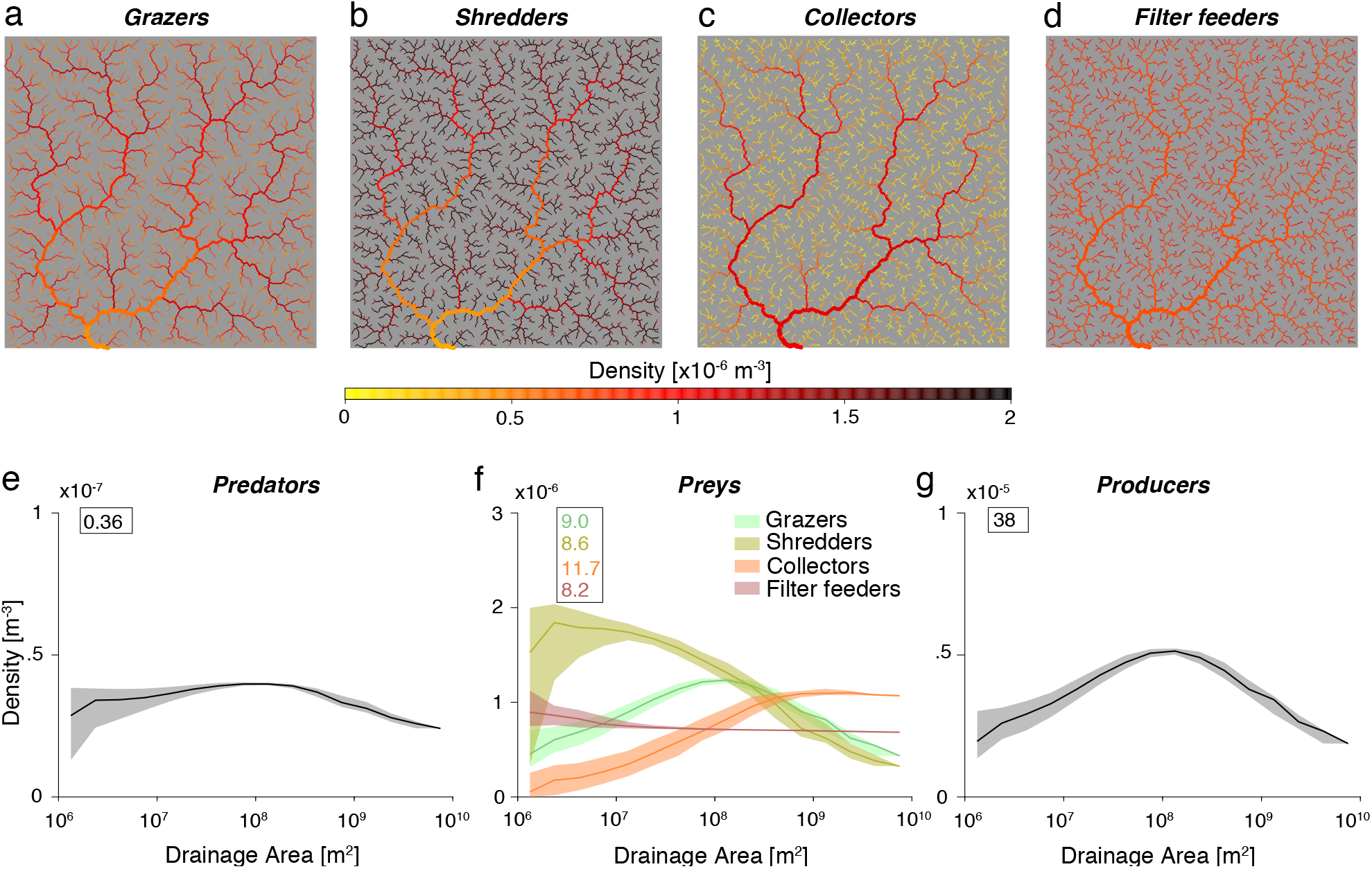
Spatial distributions of functional groups emerging from the meta-ecosystem model. a–d) Map representations of the distributions of the four main functional groups discussed in the RCC. e–g) Variation in density of functional group densities over drainage area. Solid lines represent median values calculated over reaches corresponding to each bin of Fig. 2b, while shaded areas correspond to the 2.5^th^–97.5^th^ percentile intervals evaluated across the bins of Fig. 2b. The numbers within boxes correspond to the regional biomasses of the respective functional groups (total biomass density across the whole river catchment). Note that trends in panel f correspond to the patterns shown in panels a–d.

To generalise our findings, we performed a sensitivity analysis (methodology and results are reported in the Supporting Information), where we varied two key hyper-parameters shaping food chains, namely predator feedback and pyramid top-heaviness (following Barbier and Loreau (2019)). Predator feedback expresses the ratio between consumption rates and self-regulation terms, while top-heaviness expresses the ratio between assimilation efficiency and metabolic costs. Overall, we found that the spatial patterns of consumers’ density present the same shape with respect to the default case, while regional abundances vary substantially.

We found that a linear river channel with constant width and depth does not generate spatial variations in resource and consumer densities (Fig. 4), despite the spatial effect of resource flow (see Supporting Information for a formal proof for this result). Hence, this demonstrates that the empirical spatial patterns described in the RCC can only emerge from a meta-ecosystem model that accounts for the dendritic structure of river networks. Notably, while the spatial distributions of functional groups are sensitive to the choice of landscape structure, the regional abundances tend to be similar in both landscapes. Indeed, total available habitat (i.e., total water volume) and equal total resource input from the terrestrial region were assumed equal in both landscapes. Moreover, none of the alternative landscapes built based on different assumptions on the scaling of physical variables yielded spatial patterns of functional groups similar to those postulated by the RCC (Fig. S4 – see Supporting Information for details).

**Figure 4:**
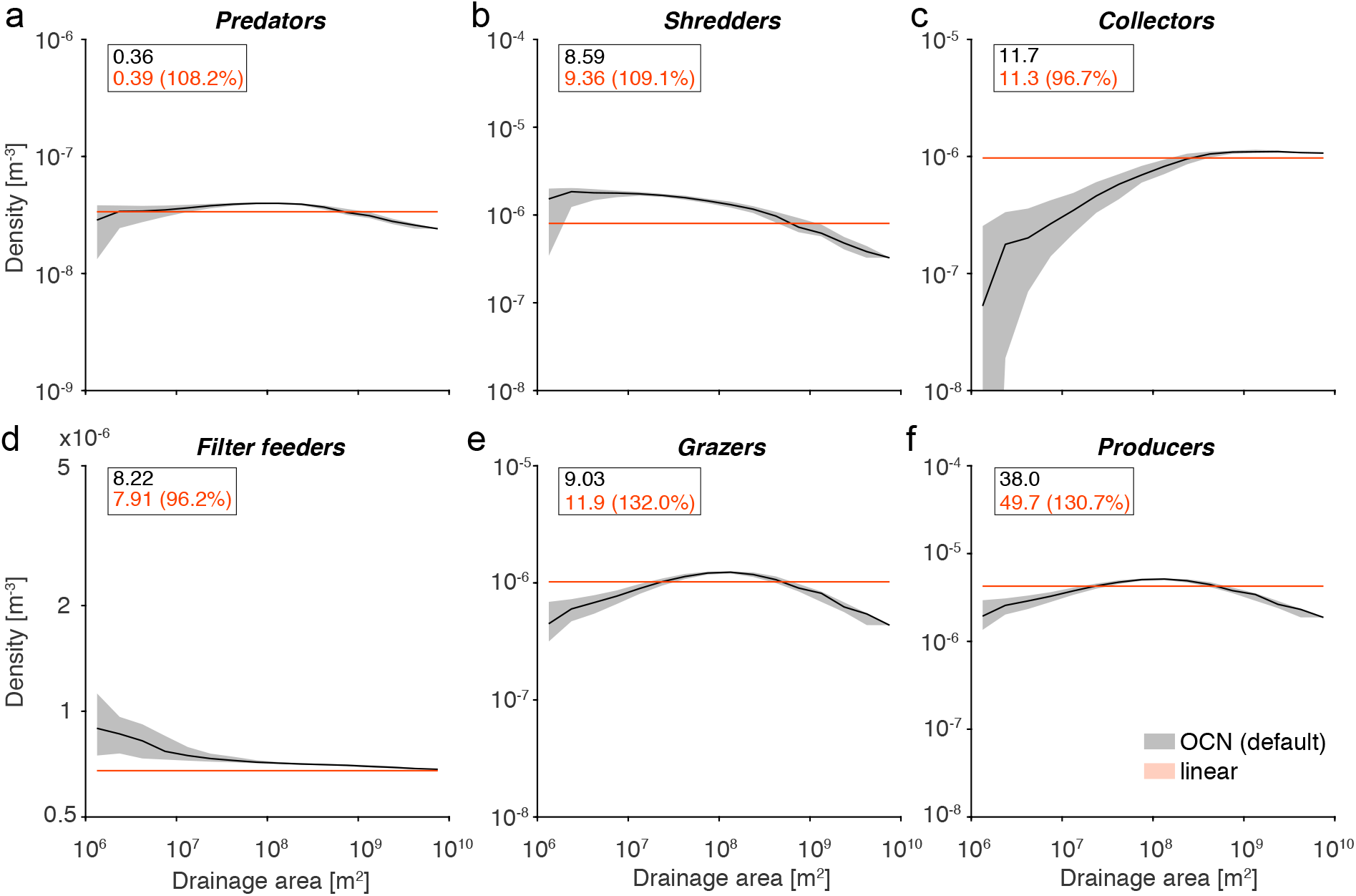
Effect of network structure on the spatial distribution and regional biomass of functional groups in stream communities. Comparison between the spatial distribution of functional groups in a linear river channel where river width and depth are constant (orange) and in a dendritic river network where river width and depth scale with drainage area (black, same as in Fig. 3). Solid lines represent median values calculated over reaches corresponding to each bin of Fig. 2b, while shaded areas correspond to the 2.5^th^–97.5^th^ percentile intervals evaluated across the bins of Fig. 2b. The numbers within boxes correspond to the regional biomasses of the respective groups; percentage values within the boxes indicate the variation in total biomass relative to the default simulation. Comparison with additional alternative landscapes based on different assumptions on scaling of physical variables is shown in Fig. S4.

Concerning the effect of variations in resource flow dynamics on riverine community patterns, the absence of hydrological transport of resources has a major effect on the spatial distribution of all functional groups in the “no flow” scenario (Fig. 5). Indeed, in this scenario, the densities of all groups decrease in the downstream direction. Given that all organisms need to rely on local terrestrial inputs in this scenario, the resulting consumer patterns essentially mirror those of input resource concentration (Fig. 2d), which tend to decrease downstream due to the dilution effect (i.e., increasing water volume downstream (Fig. 2c)). This effect is particularly strong for grazers, filterers and producers (Fig. 5c,e,f), while shredders are the least impacted. Hence, functional groups feeding on fast-flowing resources (i.e., nutrients and *DOM*) are more impacted than the ones feeding on slow-flowing resources (i.e., *CPOM*) in this scenario.

**Figure 5:**
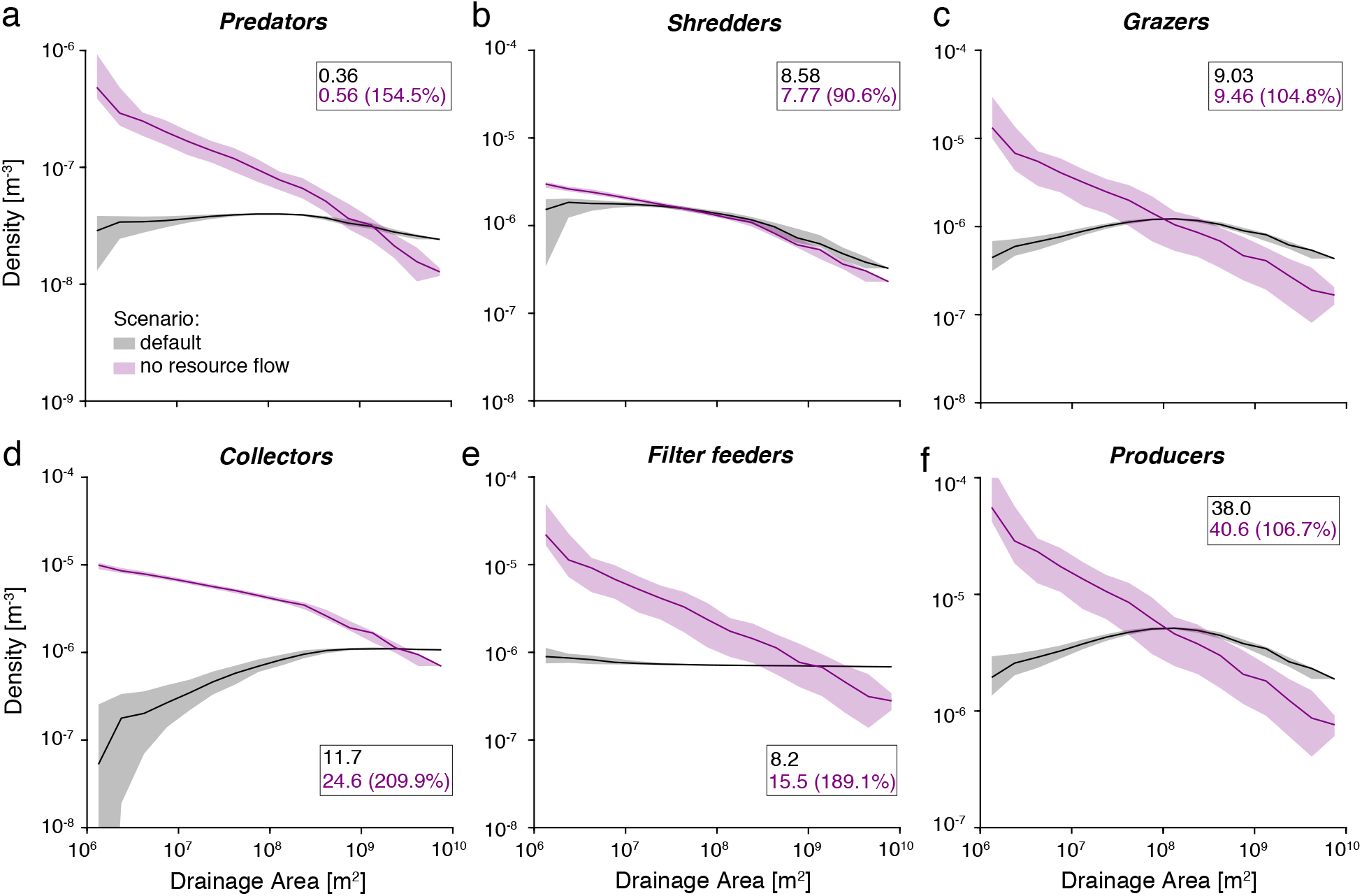
Effect of hydrological transport of resources on the spatial distribution and regional biomass of functional groups in stream communities. Comparison between the spatial distributions of functional groups with (black, same as in Fig. 3) and without (purple) hydrological transport of resources in the river network. Solid lines represent median values calculated over reaches corresponding to each bin of Fig. 2b, while shaded areas correspond to the 2.5^th^–97.5^th^ percentile intervals evaluated across the bins of Fig. 2b. The numbers within boxes correspond to the regional biomasses of the respective groups; percentage values within the boxes indicate the variation in total biomass relative to the default simulation. The corresponding resource patterns are shown in Fig. S2.

For the parameter set chosen, the regional biomasses of grazers, producers and filterers are 4.8%, 6.7% and 89.1% higher in the “no flow” scenario, respectively. For these groups, the increase in density observed in upstream reaches exceeds the decrease observed downstream. Note that this result is sensitive to the type of functional response (i.e., the relationship between a consumer’s intake rate and resource density (Holling, 1959)) used in the model (see Supporting Information for additional analyses with a type II functional response). Conversely, the regional biomass of shredders is 9.4%lower, suggesting that the decrease in *CPOM* in the most downstream reaches has a significant effect on the regional biomass of this group. The group of collectors exhibits a totally different spatial distribution in the “no flow” scenario, with very high densities in headwaters and a decreasing density in the downstream direction (Fig. 5d). The regional biomass of collectors is more than doubled, which mirrors the amount of *FPOM* available in this scenario (Fig. S2). As a result of changes in prey patterns, the spatial distribution of predator density changes as well (Fig. 5a), with higher densities in upstream reaches and a regional biomass that is 54.5% higher without hydrological transport of resources.

In the “fast” scenario, increasing the rates of hydrological transport of *CPOM* and *FPOM* has major effects on the spatial distributions and overall quantity of resources available for shredders and collectors at the regional scale (Fig. 6). The amounts of *CPOM* and *FPOM* available in the whole river network are 90.8% and 99.9% lower in this scenario, respectively (Fig. 6d,e). These changes have dramatic negative effects on the density of shredders, which is 93.6% lower at the regional scale, but shows an increasing trend in the downstream direction (Fig. 6b). We also find that shredders go extinct in 28.7% of the reaches (1058 reaches). Because *FPOM* is a by-product of shredders activity (Fig. 1), the sharp decrease in shredder density cascades to collectors, which go extinct in all reaches in this scenario (Fig. 6c). Consequently, the regional biomass of predators shrinks by more than half (Fig. 6a), with a spatial distribution following those of grazers and filterers, which are unaffected by changes in the rates of hydrological transport of *CPOM* and *FPOM*.

**Figure 6:**
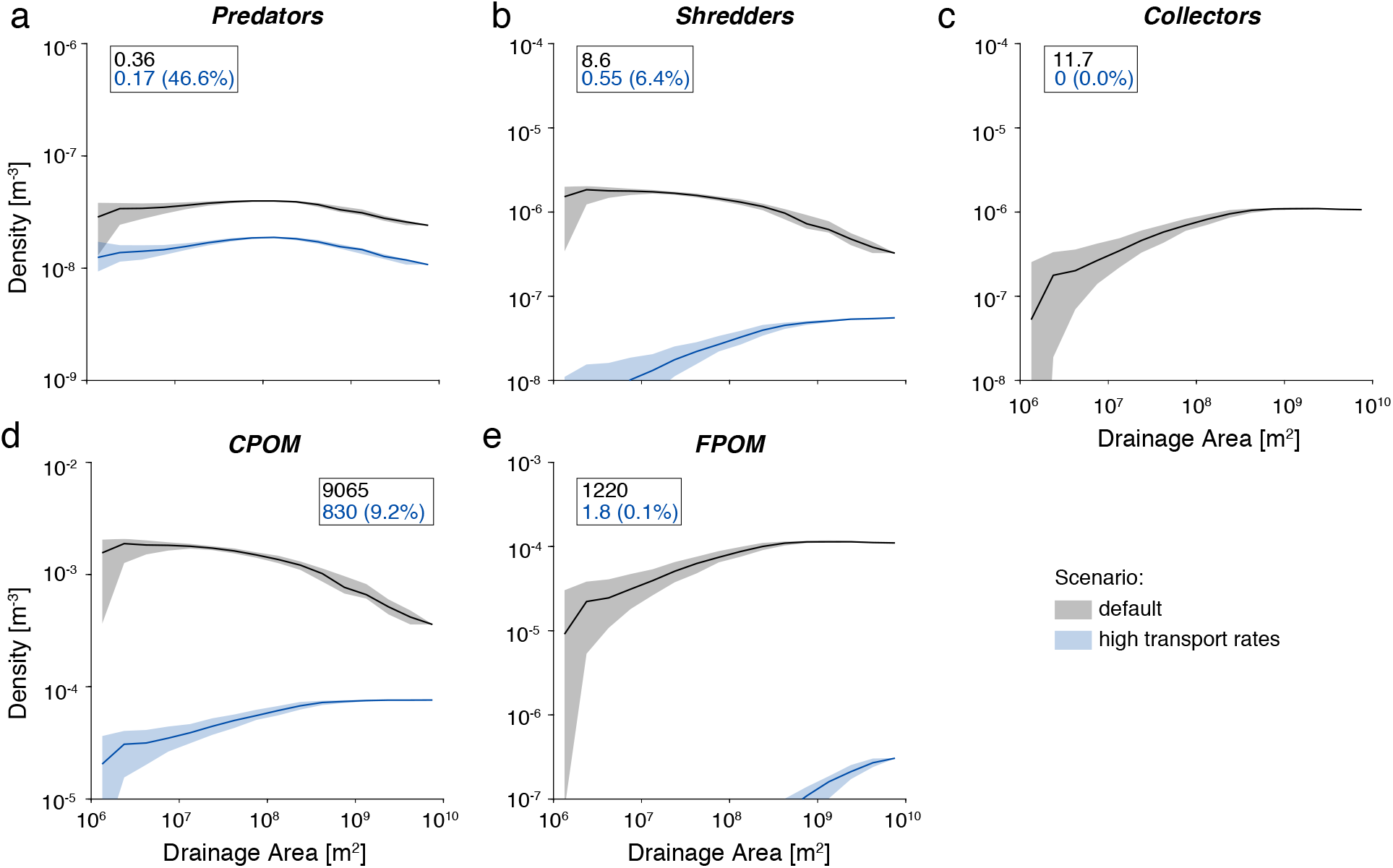
Effect of high rates of hydrological transport of resources on the spatial distribution and regional biomass of functional groups. Comparison between the spatial distributions of functional groups for the default scenario (black, same as in Fig. 3), and the scenario with high rates of hydrological transport for all resources (blue). Solid lines represent median values calculated over reaches corresponding to each bin of Fig. 2b, while shaded areas correspond to the 2.5^th^–97.5^th^ percentile intervals evaluated across the bins of Fig. 2b. The numbers within boxes correspond to the regional biomass of the respective groups; percentage values within the boxes indicate the variation in total biomass for the relative resource as compared with the default simulation.

## Discussion

By performing the first formal integration of the verbal arguments of the River Continuum Concept – a milestone that has been shaping the scientific thinking in freshwater ecology over the last four decades (Doretto et al., 2020; Vannote et al., 1980) – within a meta-ecosystem model, we showed that spatial distributions and regional biomasses of major functional groups observed in stream communities are jointly shaped by the dendritic structure and scaling attributes of river networks as well as specific rates of resource flows. Neglecting any of these aspects in modelling riverine meta-ecosystems would result in unrealistic community patterns (see also results on alternative landscapes in the Supporting Information). More generally, we showed that spatially explicit meta-ecosystem models allow understanding the interactive effects of landscape structure and resource spatial dynamics on the composition of local communities.

Model predictions did not reproduce the spatial distribution of functional groups observed in stream communities when only local ecosystem dynamics were implemented (i.e., the “no flow” scenario), which highlights the central role exerted by hydrological transport of resources in riverine ecosystems. Furthermore, the spatial distributions of functional groups were obtained without making specific assumptions on the spatial distribution of terrestrial resources (e.g., forests being more abundant at higher or inte-rmediate elevation). For instance, in our model the spatial variation of *CPOM* concentration only results from the scaling of river width and water volume with drainage area in the absence of shredders, while it is modulated by shredders’ assimilation and downstream transportation rates when shredders are present. Hence, we found that dendritic connectivity, hydrology-mediated resource flow and scaling of physical variables along a river network are sufficient to explain the widely observed community patterns postulated by the RCC. Moreover, a sensitivity analysis showed that the overall shape of these patterns is independent of the predator feedback and top-heaviness levels implemented in the food-web (see Supporting Information). Note that our work focuses on the spatial variations of functional group densities, not on the comparison of densities between functional groups in a given river reach, which would require a precise estimation of group-specific food-web parameters from empirical data, which is beyond the scope of our work.

We showed how the spatial distribution of functional groups crucially depends on the ratio between the feeding rate of each group and the rate at which their resources are transported across the river network. In particular, shredders are adapted to feed on *CPOM*, which is a slow-flowing resource. During their feeding activity, shredders produce *FPOM*, which is the resource on which collectors feed. We showed that, when the transport of *CPOM* is accelerated, which could be for instance due to the anthropogenic elimination of low-current areas or regularization of the river bed, populations of shredders are heavily depleted and this, in turn, leads to the extinction of collectors across the whole river network via a massive decrease in *FPOM* concentration. These results also support the use of metacommunity and meta-ecosystem theory for ecosystem management and restoration, as suggested by Harvey et al. (2017b), by outlining the negative effect of high rates of resource transportation in river networks on several functional groups of macroinvertebrates. The meta-ecosystem approach also formalizes the relationship between the size of the particles constituting a given resource and the rate at which it is transported via hydrological flow. This relationship is based on general physical laws and could be applied to other types of resources and other freshwater organisms, such as fishes or microbes.

Our meta-ecosystem model for river networks depends on a number of technical assumptions inherited from the RCC. First, the model does not include the dispersal of living organisms, hence making the implicit assumptions that the time scale of resource transport (which is controlled by water velocity) is much shorter than that of organismal movement across reaches, and that dispersal follows neutral dynamics, with mean distances travelled that are much smaller than the extent of the river network. In particular, these assumptions allowed us to highlight the crucial effect of resource flows on the spatial patterns of riverine communities. All these hypotheses are reasonable first approximations, yet we acknowledge that more complex dispersal dynamics can have an important role in shaping riverine metacommunities (Altermatt, 2013; Lowe and McPeek,2014; Tonkin et al., 2018) and that other freshwater organisms, such as fish, can constitute important flows of energy and nutrients, both within a river network (e.g., stream-resident fishes) and across ecosystems (e.g., potamodromous or diadromous fishes).

Second, the RCC is inspired from natural, temperate-climate rivers and the predictions of the model thus apply to natural river networks that are not subject to disconnections between river reaches due to physical barriers (e.g., dams and reservoirs) or major drying events. However, a significant fraction of river networks worldwide are fragmented (Grill et al., 2019; Belletti et al., 2020) and/or experience flow intermittence (Allen et al., 2020; Messager et al., 2021), which is likely to disrupt the longitudinal transport of resources through the network. We speculate that the effect of network fragmentation on the predictions of the RCC will depend on the location of the physical barriers or dry streams. The use of a physically-based river landscape model as done here would enable an adequate assessment of the effects of physical barriers (González-Ferreras et al., 2019) and expansions/contractions of the river network (Giezendanner et al., 2021) on stream communities.

Recent studies outlined the need for meta-ecosystem ecology to move from a very simplified and abstract representation of ecosystems to a more realistic one (Gounand et al., 2018a; Guichard, 2019). Previous theoretical studies on meta-ecosystems demonstrated that high rates of resource flow destabilize simple producer-consumer dynamics (Marleau et al., 2014). However, most studies on meta-ecosystem dynamics have focused on the effect of recycling or organism movement on ecosystem stability and productivity in small spatial networks. Here, we demonstrated that meta-ecosystem dynamics and community composition are strongly influenced by the nonrandom structure of real-world landscapes. Indeed, the spatial variation of functional groups observed in river systems only emerged from the dendritic structure and scaling of physical variables characteristic of river networks, while none of the alternative landscapes neglecting either of these aspects yielded correct predictions.

While our meta-ecosystem model was designed for river networks, the general approach proposed can be adapted to other ecological systems where meta-ecosystems dynamics are important structuring processes, such as marine shorelines, coral reef systems or estuaries (Gounand et al., 2018a; Menge et al., 2015; Spiecker et al., 2016). The application of meta-ecosystem theory to real-world landscapes crucially depends on an informed knowledge of the specific physical attributes of a landscape that constrain the direction and rates of resource flows among localities (Polis et al., 2004). Hence, a systematic quantification of cross-ecosystem flows of resources (e.g. Gounand et al. (2018b)) is central for fostering future developments of spatially explicit meta-ecosystem models and predicting the large-scale effects of human-induced changes in meta-ecosystem dynamics on biodiversity and ecosystem functioning.

## Data availability statement

The code of the meta-ecosystem for river networks is available at https://github.com/lucarraro/metaecosystem_RCC. Code will be versioned and archived on Zenodo upon acceptance.

## Acknowledgements

Funding is from the Swiss National Science Foundation Grants No PP00P3_179089 and 310030 197410 and the University of Zurich Research Priority Programme “URPP Global Change and Biodiversity” (to F.A.) and the University of Zurich Forschungskredit, grant no. K-74335-03-01 (to C.J.).

## Notes

### Competing Interest Statement

The authors have declared no competing interest.

