## Supporting Information for "Meta-ecosystem dynamics drive the spatial distribution of functional groups in river networks"

**Supplementary Figures** **15**

**Supplementary Tables** **20**

### Supplementary methods

#### Generation of the OCN

The OCN displayed in Fig. 2a was generated via the R-package *OCNet* [Carraro et al., 2020]. The OCN was constructed on a square lattice made up of 750 × 750 pixels, with each pixel's side equal to 100 m; hence, the total area spanned by the OCN equalled 5625 km$^2$. We adopted a threshold area value of 1 km$^2$ to distinguish the portion of the drainage domain effectively constituting the river network; furthermore, the maximum reach length was set to 2 km (via options *thrA* and *maxReachLength* in function *aggregate_OCN*, respectively). This resulted in a partitioning of the river network into 3688 reaches*, and a maximum Strahler order of 7. Such large dimensionality is required in order to make our results comparable with the hypotheses of the RCC (note that Vannote et al. [1980] formulated predictions for river reaches up to a Strahler order of 12, i.e. the size of the Amazon river at its mouth).

Hydraulic variables (that is, width $B$, depth $Z$ and water velocity $v$) across all network reaches were calculated with the scaling relationships of Leopold and Maddock [1953]:

$$B_i = B_o \left(\frac{A_i}{A_o}\right)^{0.5}; \qquad Z_i = Z_o \left(\frac{A_i}{A_o}\right)^{0.4}; \qquad v_i = v_o \left(\frac{A_i}{A_o}\right)^{0.1}, \tag{S1}$$

where $A_i$ is the drainage area at reach $i$; $A_o = 5625$ km$^2$ is the drainage area at the river outlet; $B_o = 24$ m, $Z_o = 5$ m and $v_o = 1.25$ ms$^{-1}$ are the (arbitrarily imposed) values of the hydraulic variables at the most downstream reach. The values of water discharge $Q$ and water volume $V$ for all reaches of the network were calculated by assuming river cross-sections as rectangular. Note that, with the assumed values for the hydraulic variables, the specific discharge at the most downstream reach is $Q_o/A_o = B_o Z_o v_o / A_o \approx 0.027$ m$^3$s$^{-1}$km$^{-2}$, a typical value for prealpine catchments [Schädler and Weingartner, 1992; Carraro et al., 2018].

#### Meta-ecosystem model

##### Resource advection

The advective terms (first terms after the equal sign) in Eqs. (1g), (1h), (1i) and (1j) were obtained as follows. The flux $\Phi_{Y,i}$ [MT$^{-1}$] of resource $Y$ exiting from a reach $i$ is equal to the mass $Y_i V_i$ of solute within that reach divided by the time that it takes for the whole mass of resource to exit the reach; such time is equal to $L_i/v_{Y,i}$ where $L_i$ is the length of reach $i$ and $v_{Y,i}$ the downstream velocity of resource $Y$ at reach $i$. We then assume that such velocity is equal to a fraction $\delta_Y$ of water velocity $v_i$. If we further assume that river

*Following Carraro et al. [2020], a reach consists in a sequence of pixels following the flow direction imposed by the OCN algorithms, which starts at a source (i.e. pixel whose drainage area is greater than or equal to *thrA*) or at the endpoint of an upstream reach, and ends at the next endpoint. Endpoints corresponds to confluences, river outlet, or pixels where the length of the upstream reach oversteps *maxReachLength*. Within a single reach $i$, hydraulic attributes such as river width $B_i$, depth $z_i$, water velocity $v_i$ and discharge $Q_i$ can be assumed constant. The portion of catchment that directly drains into reach $i$ is termed subcatchment area $A_{S,i}$. The drainage area at reach $i$ is equal to $A_i = \sum_{j\in\gamma(i)} A_{S,j}$, where $\gamma(i)$ indicates the set of reaches upstream of $i$.

cross-sections are rectangular with width $B_i$ and depth $Z_i$, it is $\Phi_{Y,i} = \delta_Y Y_i B_i Z_i v_i = \delta_Y Y_i Q_i$. The negative variation (per time unit) in resource mass at reach $i$ due to advection towards a downstream reach is $-\Phi_{Y,i}$; by assuming that water volumes do not change in time, the negative variation in resource concentration is then $-\Phi_{Y,i}/V_i$. Instead, the positive variation in resource concentration at reach $i$ due to advection from an upstream reach $j$ draining into $i$ is $\Phi_{Y,j}/V_i$.

#### Coarse particulate organic matter

*CPOM* is constituted by floating objects (leaves, wood), which however tend to get clogged easily [Wallace and Webster, 1996]. This creates accumulation of *CPOM* in specific sections of the river, prompting high local biomass of shredders due to the large resource availability. Transport of *CPOM* in rivers is thereby characterized by pulse dynamics associated to high flows. Since we refrain from including such temporal dynamics in our model, we assume that *CPOM* travels downstream at a velocity that is 1% of that of water ($\delta_{CPOM} = 0.01$).

Only a small part of the organic matter produced by forests in a given area becomes available as *CPOM* in the stream: indeed, terrestrial leaf litter can be processed by terrestrial shredders, and the byproduct of this process (or, more precisely, a fraction of it) is likely to enter the stream in the form of *DOM* transported by surface runoff and subsurface flow (see below). As a first approximation, we express the terrestrial input of *CPOM* as

$$\phi_{CPOM,i} = k_{CPOM} \min(B_i, B') L_i \,, \tag{S2}$$

where $k_{CPOM}$ expresses the amount of *CPOM* released per unit time by a unit canopy surface, while $B'$ is the maximum extent of the forest canopy above the river (that is, $B'/2$ at each bank of the reach); for reaches narrower than $B'$ (i.e., $B_i \leq B'$), the local *CPOM* input is proportional to the water surface of the reach $B_i L_i$ ($L_i$ being the reach length); for wider reaches ($B_i > B'$), it is $\phi_{CPOM} \propto B' L_i$. Eq. (S2) assumes that $k_{CPOM}$ does not depend on $i$, implying that forests are homogeneously distributed across the catchment. This hypothesis might be only partially representative of the reality, as forests are typically more abundant at intermediate altitude ranges (i.e. below the tree line and above the less steep lowlands occupied by human settlements and agricultural areas). However, it allows us to highlight the role of river connectivity and scaling of hydraulic variables in determining the spatial patterns typical of the RCC.

#### Fine particulate organic matter

Unlike *CPOM*, *FPOM* is less frequently subject to clogging and more easily transported downstream [Wallace and Webster, 1996]; its dynamics in streams have been modeled via an advection-dispersion equation with a deposition and resuspension term by Cushing et al. [1993]. Deposition velocity is of the order of magnitude of some millimeters per second, i.e. $\sim 10^3$ lower than water velocity. Moreover, *FPOM* deposited in the stream bed is not necessarily removed from the food web, as it can be consumed by benthic collectors. We thus assume that *FPOM* travels downstream at higher speed than *CPOM*, but lower speed than water: $\delta_{FPOM} = 0.5$.

We also assume that no *FPOM* input occurs from the terrestrial ecosystem, thereby implicitly hypothesizing that the terrestrial leaf litter is broken apart into particles of tiny size (*DOM*) before reaching the river via surface/subsurface flow. Hence, the local input of *FPOM* is constituted by the fraction $1 - \epsilon_S$ of in-stream *CPOM* that is broken down by shredders but not uptaken (because of "sloppy feeding" [Marks, 2019] or after excretion). It is certainly reasonable to assume that, in real settings, part of the in-stream *FPOM* enters the river network directly from the terrestrial environment (e.g. as a by-product of activity of terrestrial shredders dwelling in the proximity of a river, or by breakdown of *CPOM* operated by factors other than shredders). We here disregarded such model component for the sake of simplicity; however, we note that its inclusion would, as a first approximation, consist in a term akin to that for terrestrial input of *DOM*, and the resulting pattern of collectors would appear as a combination of the patterns of collectors and filter feeders that we found (Fig. 3).

**Dissolved organic matter and nutrients**

Because *DOM* and nutrients are made up of particles of tiny size ($\ll 1$ $\mu$m for *DOM* – molecule size for nutrients), it appears reasonable to assume that both resources are not subject to clogging and are transported downstream as passive tracers, i.e. at the same velocity as water ($\delta_{DOM} = \delta_N = 1$).

Input of terrestrial *DOM* can have multiple sources, as *DOM* can be produced both in forested and in agricultural areas [Worrall et al., 2012]. It is complicated and beyond the scope of our work to quantify the fraction of terrestrial organic matter that ultimately ends up as *DOM* in stream water. It has been observed that relevant inputs of in-stream *DOM* are due to the activity of aquatic shredders [Siders et al., 2018], although more than half of the total *DOM* in rivers is unrelated to shredders' presence [Meyer et al., 1998]; also freshwater macrophytes can generate up to 20% of *DOM* [Thomas, 1997; Reitsema et al., 2018]. Given these considerations, we here refrain from building a complete model of organic matter budget in rivers; conversely, we treat *DOM* as a different resource as opposed to *CPOM* and *FPOM*, and assume (as in Bertuzzo et al. [2017]) that the local terrestrial input of *DOM* is:

$$\phi_{DOM,i} = k_{DOM} A_{S,i} \,, \tag{S3}$$

where $A_{S,i}$ [$L^2$] is the subcatchment area and $k_{DOM}$ the flow of *DOM* released into streamflow by a unit landscape area. Analogously, and for the sake of parsimony, the local input of nutrients from the terrestrial ecosystems can be thought as constant in space and solely dependent on the local subcatchment area (as in the model of Helton et al. [2018]). Hence, it is $\phi_{N,i} = k_N A_{S,i}$.

In order to make the patterns of the different functional groups comparable, we imposed the total amounts of the three types of resources provided by the terrestrial environment (*CPOM*, *DOM*, *N*) to be all equal to 1000 (normalized mass units per day):

$$\sum_{i=1}^{n} \phi_{CPOM,i} = \sum_{i=1}^{n} \phi_{DOM,i} = \sum_{i=1}^{n} \phi_{N,i} = 1000\,\text{day}^{-1}, \tag{S4}$$

from which the coefficients $k_{CPOM}$ and $k_{DOM} = k_N$ are calculated.

**Choice of parameters for consumer-resource interactions**

The set of parameters for the default scenario was chosen in agreement with established evidence on feeding behaviour for the various functional groups, where available; alternatively, we chose parameters (as much as possible identical in value for the different functional groups) leading to an equilibrium state for system Eq. (1) in the default scenario in which state variables had non-zero values for (almost) all network reaches.

Living compartments are expressed as units of mass of organic matter per unit of volume. Note that, while the different forms of organic matter ($CPOM$, $FPOM$, $DOM$) are typically expressed in terms of carbon (although the use of the term "organic matter" rather than "organic carbon" suggests the presence of elements other than carbon [Moody and Worrall, 2017]), the nutrient ($N$) compartment essentially refers to nitrogen and phosphorus. As we are here interested in the general patterns of functional groups across river networks, we refrained from attributing empirical units to the parameters. Conceptually, in our model, assimilation rates $\alpha$ embed stoichiometric coefficients that convert units of mass between trophic levels. For the sake of convenience, masses are thus expressed in normalized units (see Table S1).

Shredders are characterized by very low assimilation efficiency [Siders et al., 2018; Marks, 2019], hence we assumed $\epsilon_S = 0.01$. On the other hand, producers are very efficient at assimilating nutrients ($\epsilon_P = 1$). For the other functional groups, in the absence of clear evidence, we set intermediate values of efficiency ($\epsilon_R = \epsilon_G = \epsilon_C = \epsilon_F = 0.1$).

Mortality rates $\mu$ were assumed to be equal to 0.01 day$^{-1}$ for all feeding groups (corresponding to an average lifespan of 100 days for all organisms), except for predators, for which $\mu_R = 0.001$ day$^{-1}$, as predicted by the metabolic theory of ecology for individuals of larger body size [Savage et al., 2004]. Feeding rates $\alpha$ for consumers at the lower trophic level of the brown food web (i.e. shredders, collectors, filterers) were set to $10^5$ m$^3$ day$^{-1}$; for producers instead, we imposed $\alpha_P = 2.5 \cdot 10^5$ m$^3$ day$^{-1}$, reflecting the higher assimilation rates of the green food web [Zou et al., 2016]. For consumers at the higher trophic level, we set higher values of the feeding rates: $\alpha_R = 10^6$ m$^3$ day$^{-1}$; $\alpha_G = 2.5 \cdot 10^6$ m$^3$ day$^{-1}$. Intra-group competition effects $\beta$ were assumed equal to $10^6$ m$^3$ day$^{-1}$ for all functional groups.

All parameters (and the respective reference values) are listed in Table S1. Parameter sets for the alternative scenarios were derived from the default set as follows: the scenario where hydrological transport of resources is neglected was obtained by setting $Q_i = 0\ \forall\, i$ in Eq. (1); the scenario with high flow rates for all resources was characterized by $\delta_{CPOM} = \delta_{FPOM} = 1$.

##### Effect of light limitation on growth of producers

In Eq. (1a), we expressed the growth rate of producers (phytoplankton) as the product between a baseline feeding rate $\alpha_P$, a light-limiting factor $l_i$ and the nutrient concentration $N_i$. As for the light-limiting factor $l_i$, we assumed

$$l_i = \frac{\overline{E_{d,i}}}{PAR}, \tag{S5}$$

where $\overline{E_{d,i}}$ is the mean downwelling irradiance along the water column of reach $i$, and $PAR$ is the irradiance of photosynthetically active radiation above the canopy cover, which, as a first approximation, we assumed constant across the whole catchment.

Light penetration in rivers can be expressed via the downwelling irradiance attenuation coefficient $K_d = -1/E_d \cdot \mathrm{d}E_d/\mathrm{d}z$ (where $E_d$ [$\mathrm{Wm}^{-2}$] is the downwelling irradiance, i.e. the quantity of solar energy that can be measured at a given river depth $z$), which was found to range between 0.2 and 5 $\mathrm{m}^{-1}$ by Davies-Colley and Nagels [2008]. The mean downwelling irradiance along the water column of reach $i$ is then:

$$\overline{E_{d,i}} = \frac{1}{Z_i}\int_0^{Z_i} E_{d,i}(z)\mathrm{d}z = E_{d0,i}\frac{1-\exp(-K_d Z_i)}{K_d Z_i} \tag{S6}$$

where $E_{d0,i}$ is the downwelling irradiance at the air-water interface of reach $i$, and $Z_i$ is the river depth. We then assumed $E_{d0,i}$ to be a fraction of $PAR$ dependent on the extent of the canopy cover above the water surface. Because in our model the forest cover was hypothesized to be homogeneous across the whole catchment, we assumed $E_{d0,i}$ to grow asymptotically towards $PAR$ as the river width $B_i$ increases:

$$\frac{E_{d0,i}}{PAR} = 1 - \exp\left(-\frac{B_i}{0.5B'}\right), \tag{S7}$$

where $B'$ is a scale width value corresponding to the maximum extent of the canopy cover above the water surface; the 0.5 coefficient at the denominator implies that the density of forests is such that a certain fraction of light can penetrate through the canopy (for instance, if $B_i = B'$, then from Eq. (S7) it is $E_{d0,i} \approx 0.86PAR$).

By combining (S5), (S6) and (S7), we obtain:

$$l_i = \frac{1-\exp(-K_d Z_i)}{K_d Z_i}\left[1 - \exp\left(-\frac{B_i}{0.5B'}\right)\right]. \tag{S8}$$

We calculated $l_i$ by imposing $K_d = 1\ \mathrm{m}^{-1}$. The distribution of values of $l_i$ as a function of drainage area is shown in Fig. 2e.

#### Search for the equilibrium state and stability check

The ordinary differential equation (ODE) system Eq. (1) is made up of 36,880 state variables (10 state variable types times 3688 network reaches). Such huge dimensionality, together with the stiffness of the ODE system, makes it computationally unfeasible to find an equilibrium point via numerical integration. Thus, we resorted to the evaluation of the equilibrium point by solving the system obtained by equalling all right-hand sides of system Eq. (1) to 0. To account for the non-linearity of the so-obtained system, we proceeded as follows: we first linearized all equations corresponding to consumers $X = \{R, G, S, C, F, P\}$ by dividing all equation terms by the respective consumer variable (i.e, discarding the solution $X = 0$); then, to solve the remaining non-linearities, we treated all consumers $X$ as parameters in the equations corresponding to resources $Y = \{CPOM, FPOM, DOM, N\}$ (in rough terms, the non-linear term $\alpha_S S_i CPOM_i$ in Eq. (1g) becomes linear if $S_i$ is treated as constant). The resulting linearized system reads:

$$\epsilon_R \alpha_R \left(G_i + S_i + C_i + F_i\right) - \mu_R - \beta_R R_i = 0\,; \tag{S9a}$$

$$\epsilon_G \alpha_G P_i - \alpha_R R_i - \mu_G - \beta_G G_i = 0\,; \tag{S9b}$$

$$\epsilon_S \alpha_S CPOM_i - \alpha_R R_i - \mu_S - \beta_S S_i = 0\,; \tag{S9c}$$

$$\epsilon_C \alpha_C FPOM_i - \alpha_R R_i - \mu_C - \beta_C C_i = 0\,; \tag{S9d}$$

$$\epsilon_F \alpha_F DOM_i - \alpha_R R_i - \mu_F - \beta_F F_i = 0\,; \tag{S9e}$$

$$\epsilon_P \alpha_P l_i N_i - \alpha_G G_i - \mu_P - \beta_P P_i \ = 0; \tag{S9f}$$

$$\frac{\delta_{CPOM}}{V_i}\left(\sum_{j=1}^{n} w_{ji} CPOM_j Q_j - CPOM_i Q_i\right) + \frac{\phi_{CPOM,i}}{V_i} - \alpha_S S_i^* CPOM_i - \lambda_{CPOM} CPOM_i = 0\,; \tag{S9g}$$

$$\frac{\delta_{FPOM}}{V_i}\left(\sum_{j=1}^{n} w_{ji} FPOM_j Q_j - FPOM_i Q_i\right) + (1-\epsilon_S)\alpha_S S_i^* CPOM_i - \alpha_C C_i^* FPOM_i - \lambda_{FPOM} FPOM_i = 0\,; \tag{S9h}$$

$$\frac{\delta_{DOM}}{V_i}\left(\sum_{j=1}^{n} w_{ji} DOM_j Q_j - DOM_i Q_i\right) + \frac{\phi_{DOM,i}}{V_i} - \alpha_F F_i^* DOM_i - \lambda_{DOM} DOM_i = 0\,; \tag{S9i}$$

$$\frac{\delta_N}{V_i}\left(\sum_{j=1}^{n} w_{ji} N_j Q_j - N_i Q_i\right) + \frac{\phi_{N,i}}{V_i} - \alpha_P l_i P_i^* N_i - \lambda_N N_i = 0\,, \tag{S9j}$$

where the asterisk marks all state variables treated as parameters. Eq. (S9) can be seen as a linear system, and can thus be written in the form $\mathbf{A}(\mathbf{x})\mathbf{x} = \mathbf{b}$, where the matrix $\mathbf{A}(\mathbf{x})$ depends on the system state. To find a solution $\mathbf{x}^{eq}$ of such system, we followed an iterative approach: starting from the initial condition $\mathbf{x}^{(0)} = \mathbf{0}$, we updated the system state via:

$$\overline{\mathbf{x}}^{(k+1)} = \mathbf{A}^{-1}\left(\mathbf{x}^{(k)}\right)\mathbf{b}, \tag{S10}$$

where $\overline{\mathbf{x}}^{(k+1)}$ represents a temporary solution at iteration $k+1$, because it might contain negative values, which hold no physical meaning. If all entries of $\overline{\mathbf{x}}^{(k+1)}$ were non-negative, the solution at iteration $k+1$

was taken as $\mathbf{x}^{(k+1)} = \overline{\mathbf{x}}^{(k+1)}$; otherwise, we solved an updated linear system:

$$\mathbf{x}^{(k+1)} = \widehat{\mathbf{A}}^{-1}\left(\mathbf{x}^{(k)}\right)\widehat{\mathbf{b}}, \tag{S11}$$

where matrix $\widehat{\mathbf{A}}\left(\mathbf{x}^{(k)}\right)$ and vector $\widehat{\mathbf{b}}$ were set equal to $\mathbf{A}\left(\mathbf{x}^{(k)}\right)$ and $\mathbf{b}$, respectively, except that all entries in the rows corresponding to the negative values of $\overline{\mathbf{x}}^{(k+1)}$ and not lying in the main diagonal of the matrix were set to 0. For instance, if collectors at a given reach $i$ were predicted to be negative by Eq. (S10), then the $i$-th row of the updated linear system Eq. (S11) would be $-\beta_C C_i = 0$, which would lead to a solution $\mathbf{x}^{(k+1)}$ in which $C_i = 0$ (i.e., replacing Eq. (S9d) for reach $i$ with $C_i = 0$). We applied Eqs. (S10) and (S11) iteratively, until a relative tolerance criterion was met:

$$\max\left(\frac{|\mathbf{x}^{(K)} - \mathbf{x}^{(K-1)}|}{\mathbf{x}^{(K)}}\right) < 10^{-6}. \tag{S12}$$

We finally set $\mathbf{x}^{eq} = \mathbf{x}^{(K)}$. We repeated the procedure herein described for all the different scenarios explored. A MATLAB script was used for this purpose.

To verify the stability of the equilibrium state $\mathbf{x}^{eq}$, we ran the full ODE system Eq. (1) via function `ode23` in MATLAB for a time span of 5,000 days. We used as initial condition a vector obtained by slightly perturbing the previously found solution $\mathbf{x}^{eq}$ by adding to each component of it a random value from a uniform distribution bound between $-10^{-3}\mathbf{x}^{eq}$ and $10^{-3}\mathbf{x}^{eq}$, plus a random value from a uniform distribution bound between 0 and $10^{-14}$; the latter term was required to perturb the null components of $\mathbf{x}^{eq}$. For all scenarios, we verified that the maximum relative error between the solution of the ODE system at the end of the time span and $\mathbf{x}^{eq}$ was below $10^{-3}$.

We also adopted a slightly different approach for equilibrium search, which exploits the fact that, in our model, state variables at a reach are only influenced by the upstream reaches, but not by the downstream ones. This makes it possible to find equilibrium states for one node at a time, starting from the headwater reaches (where the resource input from upstream is null), and subsequently moving downstream following the riverine connectivity. At a given reach, the equilibrium is found by applying the aforementioned iterative approach, except that system Eq. (S9) consists of 10 equations (given that reach $i$ is fixed), while state variables at the upstream nodes $j$ are either null (for headwater reaches) or known (from previous finding of the equilibrium state at the corresponding upstream reach). Essentially, while the original approach searches the solution of one system with 36,880 unknowns, the alternative approach searches the solution of 3688 systems with 10 unknowns each. We observed that either of the two approaches may work better in terms of computational time depending on the specific scenario setting. We hence adopted the one approach that led to fastest convergence for each scenario.

#### Model with mixed type I-type II functional responses

In our meta-ecosystem model, we assume that the feeding rate of all functional groups can be modelled with a type I functional response [Holling, 1959]. This assumption explains why we observe a very high density of producers in the upstream part of the network when we remove the hydrological transport of resources, despite the light availability being low therein (see Fig. 5e). We performed additional analyses using a type II functional response for the relationship between producers density and nutrient concentration (as in Fasham et al. [1990]). In such latter mixed model, Eqs. (1a) and (1g) were replaced, respectively, by

$$\frac{\mathrm{d}P_i}{\mathrm{d}t} = \epsilon_P \alpha'_P l_i P_i \frac{N_i}{N_i + N_0} - \alpha_G G_i P_i - \mu_P P_i - \beta_P P_i^2\,; \qquad \text{(S13a)}$$

$$\frac{\mathrm{d}N_i}{\mathrm{d}t} = \frac{\delta_N}{V_i}\left(\sum_{j=1}^{n} w_{ji} N_j Q_j - N_i Q_i\right) + \frac{\phi_{N,i}}{V_i} - \alpha'_P l_i P_i \frac{N_i}{N_i + N_0} - \lambda_N N_i\,, \qquad \text{(S13b)}$$

where $\alpha'_P$ is the predators' feeding rate for the mixed model, and $N_0 = 5 \cdot 10^{-4}$ m$^{-3}$ the half-saturation nutrient concentration. To enable comparison between the two models, we set $\alpha'_P = \alpha_P(\overline{N} + N_0)$, in which $\overline{N}$ is the mean nutrient concentration across the river network for the default simulation obtained with the original model Eq. (1). The comparison between the two models for scenarios of both presence and absence of hydrological transport of resources is shown in Fig. S3. Our results are not qualitatively modified by the implementation of the type II functional response. The only significant change occurs when the hydrological transport of resource is neglected, and consists in a lower density of grazers and producers in upstream reaches associated with lower regional biomasses (see Fig. S3). In this setting, in fact, the larger nutrient availability in the upstream reaches for the "no flow" scenario does not translate into a large resource uptake (because of a saturation effect typical of a type II functional response), and hence consumer densities are limited by the reduced light availability therein.

### Supplementary results

#### Spatial patterns of resources in the default scenario

In Fig. 2d, the input concentration of *CPOM* is characterized by a different spatial pattern with respect to those of *DOM* and *N*. Differences in spatial distributions of resource input arise because different mechanisms were hypothesized concerning the terrestrial input of the resources: while local inputs of *DOM* and *N* were assumed to be originated from the whole subcatchment area, *CPOM* input was assumed to be related to the river width: in fact, only forests that are located next to a river reach can provide *CPOM* to the meta-ecosystem, while the leaf litter originated from forests situated far from a river will be processed by terrestrial shredders, and eventually reach the river in the form of dissolved organic matter [Wallace and Webster, 1996].

The spatial patterns of resource density corresponding to the distributions of functional feeding groups of Fig. 3 are shown in Fig. S1. The spatial distributions of streamflow concentration of the different types of organic matter ($CPOM$ (Fig. S1a), $FPOM$ (Fig. S1b), $DOM$ (Fig. S1c)) tend to reproduce the density patterns of the respective consumers (shredders, collectors and filter feeders, respectively - Fig. 3f). Instead, the distribution of nutrient concentration (Fig. S1d) does not follow the pattern of producers (Fig. 3g), but rather decreases monotonically in the downstream direction; indeed, the consumption of nutrients is not solely driven by the local density of producers, but also by light availability.

Fig. S1 also shows the trend of resources in the absence of feeding groups, an alternative scenario obtained by setting to 0 all $\alpha$ parameters. Interestingly, in such case, we predict that the concentration of $CPOM$ would increase in the downstream direction (Fig. S1a), at least for drainage areas below a certain value; beyond such threshold, $CPOM$ concentration tends to decrease with increasing drainage area. The bend in instream $CPOM$ concentration is reflected by the change in the slope of the input $CPOM$ concentration (see Fig. 2d), and is explained by the fact that, beyond a certain river width (equal to $B' = 5$ m in our model, which corresponds to a drainage area of 244 km$^2$, see Eq. (S1)), the canopy cannot extend above the entire river width, hence the input of $CPOM$ per unit width decreases as drainage area (and hence width) increases (see also Eq. (S2)). Overall, in our simulation, we found that shredders consume more than 80% of the instream $CPOM$. Conversely, the patterns of $DOM$ (Fig. S1d) and nutrients (Fig. S1d) in the absence of consumers are constant in space. This is explained by the assumption of terrestrial input for both resources being only dependent on the subcatchment area (Eq. S3). For the parameter set chosen, we found that filter feeders only consume 4% of the total available $DOM$, while producers consume 17% of the total available nutrients.

#### Effect of landscape structure

To assess the effect of landscape structure and scaling of hydrological variables predicted by the OCN on spatial distribution of functional groups, we ran our model Eq. (1) on three alternative spatial settings. In all of these settings, several quantities maintained the same value as in the default (Fig. 3) scenario: total drainage area $A_o$, water discharge at the outlet $Q_o$, scaling of discharge with drainage area $Q \sim A$, number of reaches $n$, total river length $\sum L_i$, total water volume $\sum V_i$, total resource flux from the terrestrial region (equal to 1000 day$^{-1}$, see Eq. (S4)). Alternative spatial settings were designed as follows:

1. A linear river channel (without confluences) where river width and depth are constant throughout the channel. All reaches have the same length and subcatchment area (equal to $\sum L_i/n$ and $A_o/n$, respectively), and the ratio between river width and depth is fixed as $B/Z = 24/5$. Note that this scenario corresponds to the one shown in Fig. 4 of the main manuscript.

2. A linear river channel (without confluences) where river width and depth scale with drainage area according to Eq. (S1). All reaches have the same length and subcatchment area (equal to $\sum L_i/n$ and $A_o/n$, respectively), and the ratio between river width and depth at the outlet $B_o/Z_o = 24/5$ as in

the default OCN setting. $B_0$ and $Z_0$ are then recalculated such that the total water volume (i.e., total available habitat) be equal to that in the default case.

3. A river network where connectivity among reaches and reaches' lengths and subcatchment areas are the same as in the default OCN setting, but where river width and depth are constant across the network, and their ratio is $B/Z = 24/5$.

Setting 1 represents the simplest river model, made up of a linear sequence of reaches of equal size, the only spatially varying quantity being water discharge, which increases linearly in the downstream direction (because $Q \sim A$, and $A$ increases linearly in the downstream direction given that all reaches have the same subcatchment area). Setting 2 represents a case where river connectivity and distribution of drainage areas are neglected, but where scaling of hydraulic variables (and hence, changes in reach size in the downstream direction) are considered. Finally, setting 3 preserves the correct network topology but disregards spatial variations in reach size. Unless otherwise specified, all parameter values were taken from Table S1.

Fig. S4 shows the effect of alternative spatial settings on spatial patterns of resource and consumer densities for the different. Overall, these patterns tend to differ substantially with respect to the default OCN case. Remarkably, the linear setting without scaling of hydraulic variables leads to constant patterns of consumer and resource density. A formal proof for this fact is provided in the following paragraph. In the case of the linear channel with scaling of hydraulic variables, densities of all functional groups (excluding filter feeders) tend to be much lower than in the default case. In particular, shredders' density (Fig. S4b) tends to peak for intermediate values of drainage area, and moderately decline further downstream. Collectors' density (Fig. S4c) is null in the 2% most upstream reaches, and increases downstream (with values much lower than those predicted for the default case, except towards the outlet, where values tend to be more similar). Peak densities of grazers (Fig. S4e) and producers (Fig. S4f) are localized closer to the outlet than it was the case for the default setting. As a result, in this setting predators' density (Fig. S4a) is predicted to increase in the downstream direction.

Finally, the network setting without scaling of hydraulic variables leads to patterns of consumers that, in most cases, tend to decrease in the downstream direction (especially for predators, shredders, grazers and producers), while collectors' density appears highly variable in the upstream part of the network and tends to the density observed in the default case towards the outlet. In all three settings, filter feeders' density (Fig. S4d) is spatially constant and very similar to that of the default case; indeed, both the input of *DOM* in the river and the consumption process by this functional group were assumed to be independent of any reach-specific variable (such as river width, which is key for *CPOM* input and thus shredder density, and depth, which influences producers' assimilation rate via the light limitation factor).

It is worthwhile to note that, while spatial patterns of consumers and resources highly depend on the choice of spatial setting, the regional abundances tend to be similar to those of the default case. Indeed, these comparisons were performed by imposing equal total available habitat (i.e., total water volume), equal rate of removal of resources from the system (which is controlled by the discharge at the outlet

$Q_o$), and equal total resource input from the terrestrial region. Hence, the main differences among spatial settings lie in the different distributions of drainage areas across reaches and in the consequent different distributions of hydraulic variables, and in particular of water volume (Fig. S5). While network settings have a disproportionate number of headwaters (and hence of reaches with low drainage area) with respect to downstream reaches, linear settings have a single headwater, with drainage area increasing linearly in the downstream direction, hence the distributions along the $x$-axes of Fig. S5 are shifted towards the right. This explains why for instance, despite the collector density in the linear setting with scaling being an order of magnitude lower than that in the default case for most drainage area values, the regional abundance for this functional group is more than 80% of that in the default case (Fig. S4c). Indeed, most of reaches in the linear setting with scaling have high drainage area values, where the predicted collector density is similar to that of the default case.

The fact that large variations in the consumer patterns depending on the setting type are found for low drainage area values (consider e.g. shredders, Fig. S4b) essentially depends on the distribution of water velocity values, which was not preserved in the formulation of alternative settings. Indeed, in our model, water velocity at a reach can be expressed as $QL/V$; while $Q$ values for a given drainage area were fixed across settings by hypothesis (i.e., because $Q_o$ and $Q \sim A$ are preserved across setting types), and reach length varies independently of drainage area, the distribution of water volumes varies substantially with $A$ (Fig. S5). In the default OCN case, water volumes increase in the downstream direction because of the scaling of width and depth with drainage area; the variability in water volumes for a given drainage area is due to differences in reach lengths (this latter feature is also observed in the network-without-scaling setting, while reach lengths are equal by construction in the linear settings). Water volume values for low drainage ares in the OCN are higher than in the linear-with-scaling setting, but lower than in the two no-scaling settings, which results in water velocities upstream being highest in the linear-with-scaling setting, and lowest for the no-scaling settings. This explains why the pattern of shredder density increases downstream in the linear-with-scaling setting (Fig. S4b); in analogy with the scenario of fast resource transportation, in this setting shredders upstream consume a reduced (with respect to the default case) amount of $CPOM$ because this resource travels faster downstream. The opposite is true for the network-without-scaling setting.

**State variables are spatially constant in linear, non-scaling landscape**

*Proof.* For the sake of simplicity, let's consider a model constituted by a single resource flowing downstream and its respective consumer (say, nutrients $N$ and producers $P$). Let the most upstream reach (termed 1) have equilibrium nutrient concentration and producer density $N_1^*$ and $P_1^*$, respectively. From Eq. (1a), it is $P_1^* = \left(\epsilon_P \alpha_P l N_1^* - \mu_P\right) / \beta_P$. Note that the light factor $l$ appears without subscript as it is spatially constant in this setting. Moreover, from Eq. (1g), it is:

$$\alpha_P l V P_1^* N_1^* = \phi_N - \delta_N Q_1 N_1^* - \lambda_N V N_1^* \,, \tag{S14}$$

where again all subscripts referred to reach-specific quantities have been removed. At the next downstream reach (identified with subscript 2), let $N_2^*$ and $P_2^* = (\epsilon_P \alpha_P l N_2^* - \mu_P)/\beta_P$ be the equilibrium nutrient concentration and producer density therein, respectively. From Eq. (1g), it is:

$$N_2^* = \frac{\phi_N + \delta_N Q_1 N_1^*}{\delta_N Q_2 + \alpha_P l V P_2^* + \lambda_N V}\,. \tag{S15}$$

Subtracting $N_1^*$ from both sides of Eq. (S15), and considering that $Q_2 = 2Q_1$ in this setting, one gathers

$$N_2^* - N_1^* = \frac{\phi_N - \delta_N Q_1 N_1^* - \lambda_N V Q_1 N_1^* - \alpha_P l V P_2^*}{2\delta_N Q_1 + \alpha_P l V P_2^* + \lambda_N V}\,. \tag{S16}$$

Substituting Eq. (S14) in Eq. (S16), and recalling the definitions of $P^*$ expressed as a function of $N^*$, one gathers

$$N_2^* - N_1^* = \frac{\epsilon_P \alpha_P^2 l^2 V}{\beta_P \left(2\delta_N Q_1 + \alpha_P l V P_2^* + \lambda_N V\right)}\left(N_1^* - N_2^*\right). \tag{S17}$$

Given that the fraction in the r.h.s. of Eq. (S17) is positive by construction, equality Eq. (S17) can only be true if $N_1^* = N_2^*$ and hence also $P_1^* = P_2^*$. The same reasoning can be applied to any other pair of reaches (e.g., 1 and 3, where 3 is the reach downstream of 2, where $Q_3 = 3Q_1$). □

#### Sensitivity analysis

Given the large number of parameters related to the feeding functional groups of our model (see Table S1), we refrained from running a full sensitivity analysis. Instead, following Barbier and Loreau [2019], we focused our attention on two hyper-parameters shaping food chains, namely the predator feedback $\lambda$ and the top-heaviness $\kappa$. Predator feedback $\lambda$ expresses the ratio between consumption rates and self-regulation terms, and identifies whether the food chain has a pyramidal (low $\lambda$) or cascading (high $\lambda$) shape; in the context of our model, it is $\lambda \approx \alpha/\beta$. Top-heaviness $\kappa$ controls whether the pyramidal food chain has a top-down (high $\kappa$) or bottom-up (low $\kappa$) shape. Top-heaviness expresses the ratio between assimilation efficiency and metabolic costs; in the context of our model, given that we disregarded differences in metabolic costs among the different trophic levels, it is $\kappa \approx \epsilon$.

We explored four scenarios in which we ran model simulations by altering groups of parameters corresponding to higher/lower predator feedback and higher/lower top-heaviness, respectively:

1. In the high $\lambda$ scenario, we multiplied all feeding rates $\alpha$ of Table S1 by 3. Importantly, we noted that further increasing $\alpha$'s with respect to these values led to non-equilibrium in some network reaches. While investigation of non-equilibrium dynamics in our model is certainly of interest, this aspect was deemed out of scope with respect to the focus of this work.

2. In the low $\lambda$ scenario, we divided all feeding rates $\alpha$ of Table S1 by 3.

3. In the high $\kappa$ scenario, we multiplied all feeding efficiencies $\epsilon$ (barring $\epsilon_P$) of Table S1 by 10. We maintained $\epsilon_P = 1$ as in the default case, since efficiencies cannot physically exceed unity.

4. In the low $\kappa$ scenario, we divided all feeding efficiencies $\epsilon$ (barring $\epsilon_P$) of Table S1 by 10.

Unless otherwise stated, all parameters maintained their default values from Table S1.

Results for the sensitivity analysis are presented in Fig. S6. Overall, the spatial patterns of consumers' density present the same shape with respect to the default case, while regional abundances vary substantially. In general, scenarios with either high predator feedback or high top-heaviness tend to result in higher regional abundances for most feeding groups, while the opposite is true for scenarios with either low $\lambda$ or low $\kappa$. An interesting exception to this trend is constituted by producers (Fig. S6f), for which predicted regional abundance in the default setting is the highest compared to the four alternative scenarios. A likely explanation is that, for the high $\lambda$ scenarios, a cascading-like pattern can be observed (because the grazer-producer food chain is the longest in our model), according to which grazers increase their density as compared to the default scenario (Fig. S6e), while producers do not. As for the high $\kappa$ scenario, the increase in grazers' density (with respect to the default case) coupled with a decrease in producers' density could be a hint of a top-down-shaped pyramid; however, note also that the efficiency of producers was not changed with respect to the default case.

In the high $\lambda$ scenario, peaks in grazer and producer densities tend to occur a bit more upstream than for the default case, and descending limbs in the downstream part of the catchment tend to be steeper (see e.g. shredders or collectors). This is caused by higher resource depletion occurring in this scenario. Conversely, in the low $\kappa$ scenario, predators (Fig. S6a) are unable to survive in any reach of the network; moreover, grazers are predicted to be absent in 43% of the network reaches, located in both the most upstream and downstream portions of the network.

### Supplementary Figures

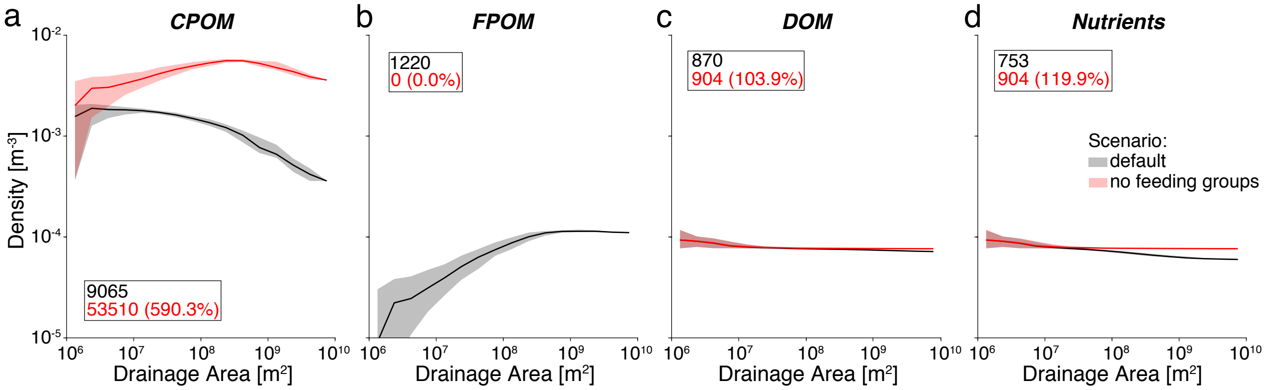

Figure S1: Effect of biotic processes on the spatial variation of resource concentration along the river network. Black color refers to the default simulation, i.e. corresponding to the spatial distribution of feeding groups shown in Fig. 3; and red to a scenario where all feeding groups are absent. Plot construction is as in Fig. 4.

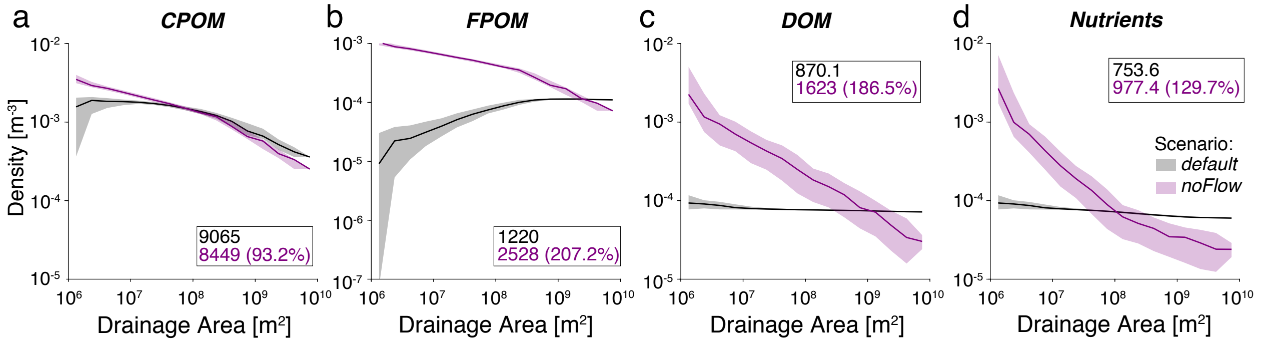

Figure S2: Effect of hydrological transport of resources on the structure of riverine food webs. Comparison between the spatial distribution of resources in the presence (black lines, corresponding to the trends shown in Fig. S1) and in the absence of streamflow (purple). Plot construction is as in Fig. 4. The related comparison of patterns of feeding groups is shown in Fig. 4.

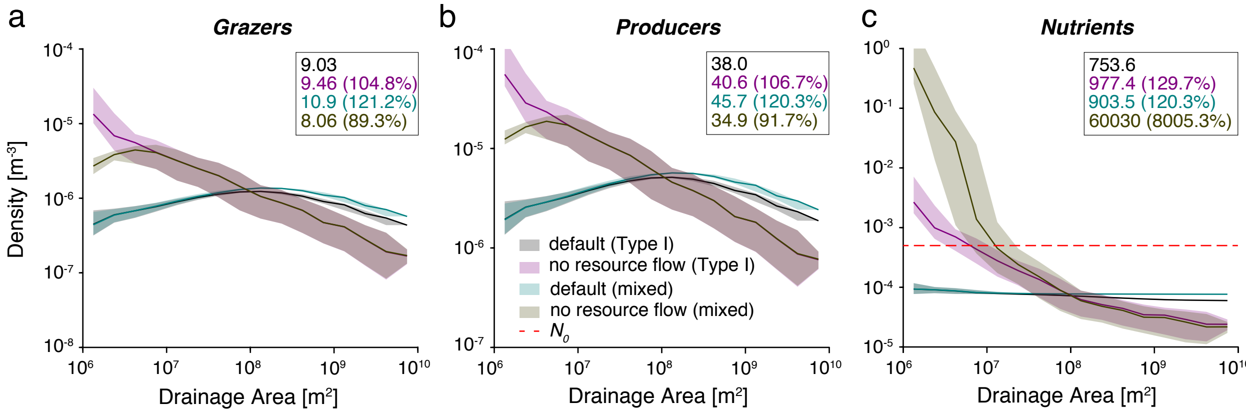

Figure S3: Effect of assumption of a type II functional response between producers and nutrients for scenarios of both presence and absence of hydrological transport of resources. Black and purple curves are the same as in Fig. 4. Plot construction is as in Fig. 4.

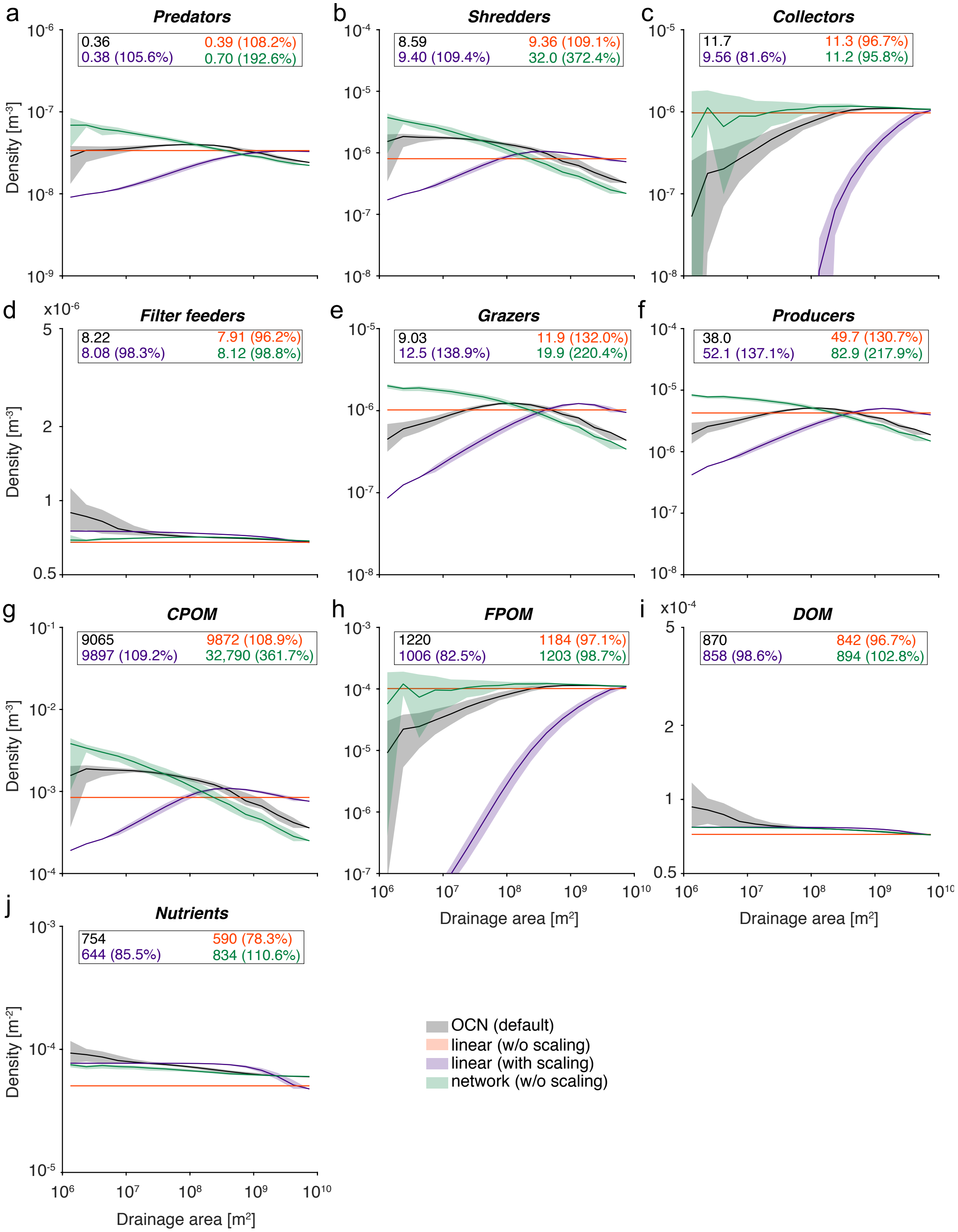

Figure S4: Effect of landscape structure on the structure of riverine food webs. Orange lines and shades correspond to results for the linear river channel landscape shown in Fig. 4. Plot construction is as in Fig. 4.

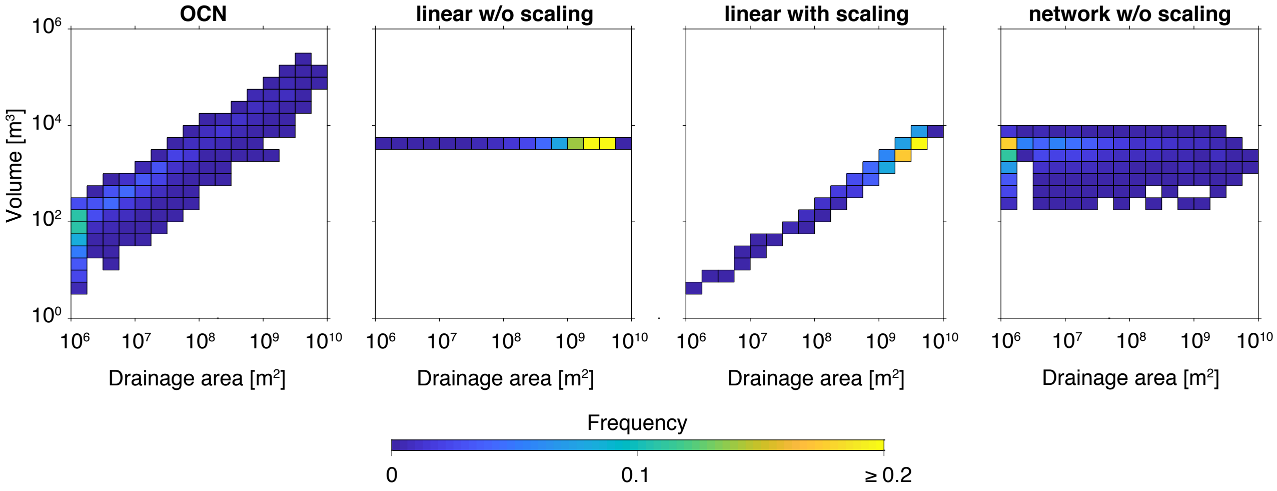

Figure S5: 2D-histograms of drainage area and water volume values for the four different spatial settings analyzed.

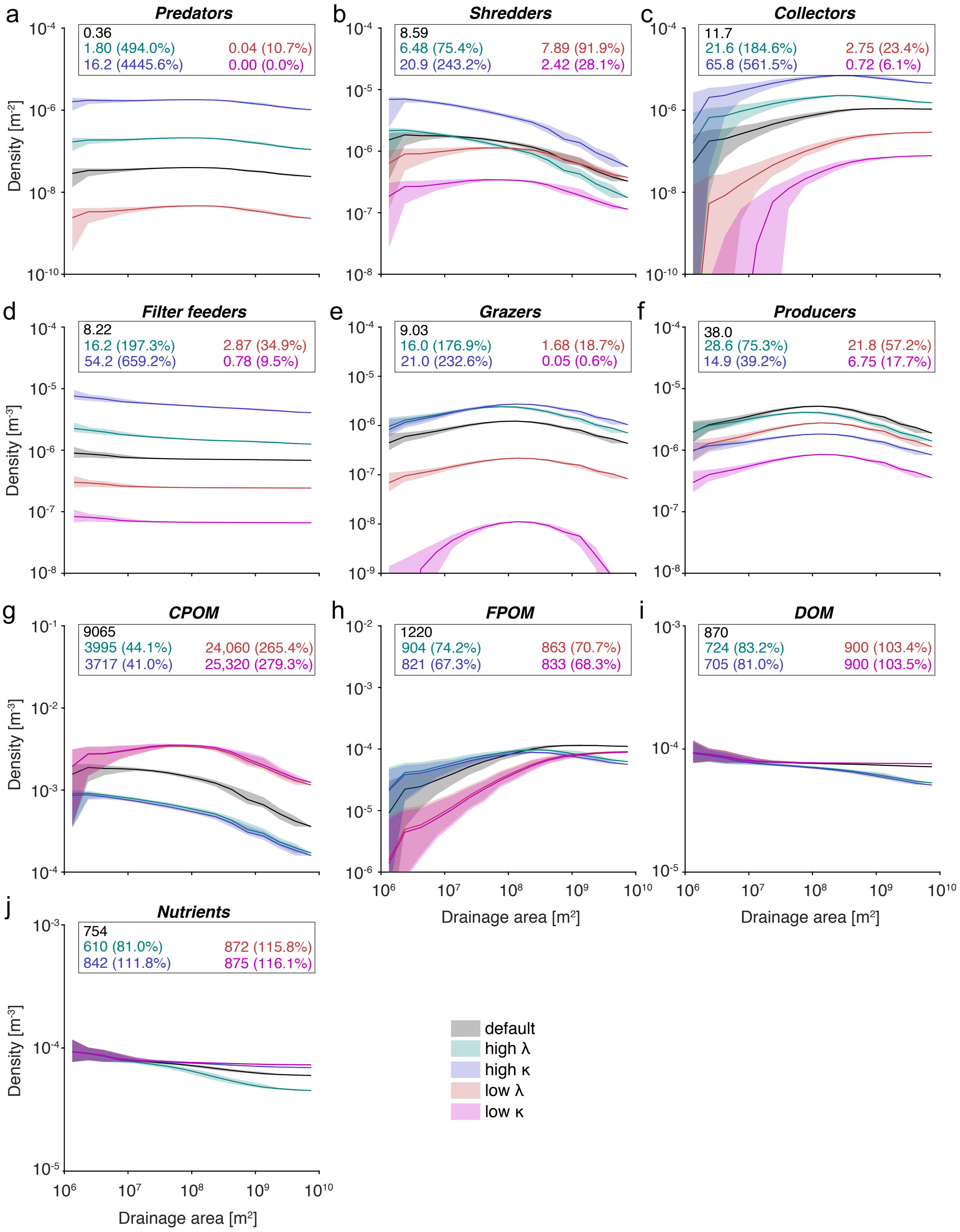

Figure S6: Assessment of sensitivity of model results to variations in food-web parameters.

403 # Supplementary Tables

Table S1: List of parameters for the default scenario. Note that all state variables of model Eq. (1) are expressed as densities [$ML^{-3}$], with masses expressed in normalized units. Parameter values are discussed in section "Choice of parameters for consumer-resource interactions" of the Supporting Information.

| Symbol | Description | Dimension* | Reference value |
|---|---|---|---|
| **Parameters related to the feeding functional groups** | | | |
| $\alpha_R$ | Predators' feeding rate | $L^3M^{-1}T^{-1}$ | $10^6$ m$^3$ day$^{-1}$ |
| $\alpha_G$ | Grazers' feeding rate | $L^3M^{-1}T^{-1}$ | $2.5 \cdot 10^6$ m$^3$ day$^{-1}$ |
| $\alpha_S$ | Shredders' feeding rate | $L^3M^{-1}T^{-1}$ | $10^5$ m$^3$day$^{-1}$ |
| $\alpha_C$ | Collectors' feeding rate | $L^3M^{-1}T^{-1}$ | $10^5$ m$^3$day$^{-1}$ |
| $\alpha_F$ | Filterers' feeding rate | $L^3M^{-1}T^{-1}$ | $10^5$ m$^3$day$^{-1}$ |
| $\alpha_P$ | Producers' feeding rate | $L^3M^{-1}T^{-1}$ | $2.5 \cdot 10^5$ m$^3$day$^{-1}$ |
| $\beta_R$ | Predators' intragroup | $L^3M^{-1}T^{-1}$ | $10^6$ m$^3$ day$^{-1}$ |
| $\beta_G$ | Grazers' intragroup | $L^3M^{-1}T^{-1}$ | $10^6$ m$^3$ day$^{-1}$ |
| $\beta_S$ | Shredders' intragroup competition effect | $L^3M^{-1}T^{-1}$ | $10^6$ m$^3$ day$^{-1}$ |
| $\beta_C$ | Collectors' intragroup competition effect | $L^3M^{-1}T^{-1}$ | $10^6$ m$^3$ day$^{-1}$ |
| $\beta_F$ | Filterers' intragroup competition effect | $L^3M^{-1}T^{-1}$ | $10^6$ m$^3$ day$^{-1}$ |
| $\beta_P$ | Producers' intragroup competition effect | $L^3M^{-1}T^{-1}$ | $10^6$ m$^3$ day$^{-1}$ |
| $\mu_R$ | Predators' mortality rate | $T^{-1}$ | 0.001 day$^{-1}$ |
| $\mu_G$ | Grazers' mortality rate | $T^{-1}$ | 0.01 day$^{-1}$ |
| $\mu_S$ | Shredders' mortality rate | $T^{-1}$ | 0.01 day$^{-1}$ |
| $\mu_C$ | Collectors' mortality rate | $T^{-1}$ | 0.01 day$^{-1}$ |
| $\mu_F$ | Filterers' mortality rate | $T^{-1}$ | 0.01 day$^{-1}$ |
| $\mu_P$ | Predators' mortality rate | $T^{-1}$ | 0.01 day$^{-1}$ |
| $\epsilon_R$ | Predators' assimilation efficiency | - | 0.01 |
| $\epsilon_G$ | Grazers' assimilation efficiency | - | 0.1 |
| $\epsilon_S$ | Shredders' assimilation efficiency | - | 0.01 |
| $\epsilon_C$ | Collectors' assimilation efficiency | - | 0.1 |
| $\epsilon_F$ | Filterers' assimilation efficiency | - | 0.1 |
| $\epsilon_P$ | Producers' assimilation efficiency | - | 1 |
| **Parameters related to the resources** | | | |
| $\delta_{CPOM}$ | Relative downstream velocity of $CPOM$ | - | 0.01 |
| $\delta_{FPOM}$ | Relative downstream velocity of $FPOM$ | - | 0.5 |
| $\delta_{DOM}$ | Relative downstream velocity of $DOM$ | - | 1 |
| $\delta_N$ | Relative downstream velocity of nutrients | - | 1 |
| $\lambda_{CPOM}$ | Loss rate of $CPOM$ | $T^{-1}$ | 0.01 day$^{-1}$ |
| $\lambda_{FPOM}$ | Loss rate of $FPOM$ | $T^{-1}$ | 0.01 day$^{-1}$ |
| $\lambda_{DOM}$ | Loss rate of $DOM$ | $T^{-1}$ | 0.01 day$^{-1}$ |
| $\lambda_N$ | Loss rate of nutrients | $T^{-1}$ | 0.01 day$^{-1}$ |
| $k_{CPOM}$ | Rate of $CPOM$ biomass release | $MT^{-1}L^{-2}$ | From Eq. (S4) |

Table S1: List of parameters for the default scenario. Note that all state variables of model Eq. (1) are expressed as densities [$ML^{-3}$], with masses expressed in normalized units. Parameter values are discussed in section "Choice of parameters for consumer-resource interactions" of the Supporting Information.

| Symbol | Description | Dimension* | Reference value |
|---|---|---|---|
| $k_{DOM}$ | Rate of *DOM* biomass release | $MT^{-1}L^{-2}$ | From Eq. (S4) |
| **Landscape parameters** | | | |
| $A_{S,i}$ | Subcatchment area | $L^2$ | Given by OCN |
| $B_i$ | River width in reach $i$ | L | Given by OCN |
| $B'$ | Maximum extent of canopy cover above river | L | 5 m |
| $L_i$ | Reach length | L | Given by OCN |
| $l_i$ | Ligth factor | - | Eq. (S8) |
| $Q_i$ | Water discharge in reach $i$ | $L^3T^{-1}$ | Given by OCN |
| $V_i$ | Water volume of reach $i$ | $L^3T^{-1}$ | Given by OCN |
| $Z_i$ | Water depth of reach $i$ | L | Given by OCN |
| $w_{ji}$ | Entry of adjacency matrix: $w_{ji} = 1$ if reach $j \to i$; $w_{ji} = 0$ otherwise | - | Given by OCN |
| $A_i$ | Drainage area at reach $i$ | $L^2$ | |

*Mass is expressed in normalized units.
